# AlphaFold3 in Drug Discovery: A Comprehensive Assessment of Capabilities, Limitations, and Applications

**DOI:** 10.1101/2025.04.07.647682

**Authors:** Haiyang Zheng, Hanfeng Lin, Adebowale A. Alade, Jingjing Chen, Erika Y. Monroy, Min Zhang, Jin Wang

**Affiliations:** Verna and Marrs McLean Department of Biochemistry and Molecular Pharmacology, Baylor College of Medicine, Houston, Texas 77030, United States; Center for NextGen Therapeutics, Baylor College of Medicine, Houston, Texas 77030, United States; Department of Molecular and Cellular Biology, Baylor College of Medicine, Houston, Texas 77030, United States

## Abstract

Accurate prediction of protein-ligand interactions remains a cornerstone challenge in drug discovery. AlphaFold3 (AF3), a recent breakthrough Diffusion Transformer model, holds significant promise for structural biology, but its performance across diverse pharmaceutical applications requires systematic evaluation. In this study, we comprehensively benchmark AF3’s capabilities using carefully curated datasets, examining its performance in binary protein-ligand complexes, apo/holo structural variations, GPCR-ligand conformations, ternary systems, and inhibitor affinity prediction.

Our analysis reveals that AF3 excels at predicting static protein-ligand interactions with minimal conformational changes, significantly outperforming traditional docking methods in side-chain orientation accuracy. However, we identify critical limitations: AF3 struggles with protein-ligand complexes involving significant conformational changes (>5Å RMSD), demonstrates a persistent bias toward active GPCR conformations regardless of ligand type, performs poorly on ternary complex prediction, and lacks reliable affinity ranking capability. Notably, AF3’s performance declined significantly on structures released after its training cutoff date, suggesting potential memorization rather than physical understanding of molecular interactions.

We explored AF3’s practical utility through applications in chemoproteomics data interpretation, drug resistance mutation prediction, and kinome profiling simulation. AF3 demonstrated value as a “true-hit binary interaction modeler,” capable of generating reliable structural models for experimentally validated binding pairs. However, its ranking metrics showed minimal correlation with experimental binding affinities and limited ability to differentiate across the kinome, highlighting the need for integration with physics-based scoring methods.

Our findings indicate that while AF3 represents a significant advancement in protein-ligand structure prediction, it requires complementary approaches to address its limitations in conformational sampling, affinity ranking, and complex system modeling. Recent developments like YDS Ternoplex suggest that enhanced sampling techniques can overcome some of these limitations. The optimal strategy for leveraging AF3 in drug discovery likely involves its integration into hybrid computational pipelines that combine AI-based prediction with physics-based refinement and experimental validation.

## Introduction

The accurate prediction of protein-ligand interactions represents a fundamental challenge at the intersection of structural biology and drug discovery. These molecular recognition events underpin virtually all biological processes and serve as the mechanistic foundation for therapeutic intervention. Despite decades of methodological advancement, achieving high-fidelity predictions of these complex interactions has remained elusive due to the inherent complexity of biomolecular systems, involving intricate networks of weak non-covalent interactions, conformational dynamics, and solvation effects.

Traditional computational approaches to protein-ligand interaction prediction, primarily relying on molecular docking algorithms, have provided valuable insights but suffer from significant limitations. While these methods can generate reasonable binding poses in many cases, they frequently struggle with accurate prediction of side-chain orientations—critical determinants of binding specificity and affinity.^1^ This inaccuracy often results in low confidence predictions that can mislead structure-activity relationship (SAR) efforts in medicinal chemistry campaigns. Furthermore, experimental structure determination of protein-ligand complexes typically requires weeks or months of dedicated effort, creating a substantial bottleneck in the drug discovery pipeline.^2^

The emergence of machine learning and artificial intelligence has catalyzed a paradigm shift in structural biology. AlphaFold2 (AF2), introduced in 2021, demonstrated unprecedented accuracy in protein structure prediction, effectively solving a 50-year scientific challenge and transforming the field.^3^ This breakthrough spurred the development of numerous derivative tools and approaches based on the underlying principles of AF2.^4,5^ Building upon this foundation, DeepMind recently unveiled AlphaFold3 (AF3), representing a significant architectural evolution with its Diffusion Transformer design. According to DeepMind’s internal benchmarking, AF3 exhibits remarkable accuracy in predicting protein-ligand interactions, potentially addressing a critical gap in computational drug discovery.^6^

However, the practical utility of AF3 across diverse drug discovery scenarios remains incompletely characterized. While initial demonstrations have highlighted its capabilities, a systematic evaluation of its performance boundaries and limitations in real-world pharmaceutical applications is essential for its effective integration into drug development workflows. Critical questions persist regarding AF3’s ability to predict induced conformational changes, accommodate diverse binding modes, distinguish between active and inactive compounds, and model complex multi-component systems relevant to emerging therapeutic modalities.

This study aims to conduct a comprehensive assessment of AF3’s performance across a spectrum of scenarios directly relevant to contemporary drug discovery challenges. Our evaluation encompasses several distinct categories: (1) binary protein-ligand interactions, with particular attention to both static complexes and those involving significant conformational change; (2) protein conformational states induced by ligands, focusing on G protein-coupled receptors (GPCRs) and the distinction between agonist and antagonist binding; (3) ternary protein-ligand systems critical for molecular glue and PROTAC development; and (4) inhibitor affinity prediction capability essential for compound ranking and prioritization.

Using carefully curated datasets including PLINDER^7^ and GPCR-DB^8^, we systematically evaluate AF3’s structural prediction fidelity, interaction recovery rates, and ability to differentiate compounds based on binding affinity. Furthermore, we explore practical applications of AF3 in addressing real-world drug discovery challenges, including the structural interpretation of chemoproteomics screening data, prediction of drug resistance mutations, and simulation of kinome selectivity profiles.

Through this comprehensive assessment, we aim to delineate both the remarkable capabilities and inherent limitations of AF3, providing the scientific community with a nuanced understanding of its optimal application domains within drug discovery workflows. By identifying specific performance boundaries, we establish a foundation for the strategic integration of AF3 with complementary computational and experimental approaches, potentially accelerating the discovery and optimization of novel therapeutic agents.

## Results

### Evaluation: Protein-Ligand Binary Interactions - Foundations and Performance Limitations

The binary protein-ligand interaction represents the fundamental molecular recognition process underlying drug action. In this study, we conducted a comprehensive assessment of AlphaFold3 (AF3) and Chai-1, two state-of-the-art AI prediction models, on this critical task. Chai-1, developed by Chai Discovery, offers similar capabilities to AF3 with a unified architecture for predicting diverse biomolecular structures including proteins, small molecules, nucleic acids, and covalent modifications.^9^

Previous research has indicated that AI prediction models often struggle with accurately capturing conformational changes induced by ligand binding.^10^ To systematically investigate this limitation, we strategically divided our benchmark into two distinct categories: static complexes (protein-ligand structures where the protein conformation closely resembles the unbound apo state, RMSD < 0.5Å) and dynamic complexes (structures exhibiting significant conformational rearrangements upon ligand binding, RMSD > 5Å). Additionally, we implemented a temporal segregation of test cases based on training cutoff dates to assess potential memorization effects in these models.

For our experimental evaluation, we randomly selected 150 structures from each category and generated predictions using both AF3 and Chai-1. When comparing predicted structures to experimental references from the PDB, we observed several noteworthy trends (Figure 1). Without training cutoff filtration, both models demonstrated markedly superior performance on Static Complexes compared to Dynamic Complexes, confirming our hypothesis regarding conformational adaptability limitations (Figure 1A and B). Interestingly, this performance gap substantially diminished for structures released after the training cutoff date (Figure 1C and D). Comparative analysis across temporal divisions revealed a general decline in predictive accuracy for post-cutoff structures (Figure 2).

**Figure 1.**
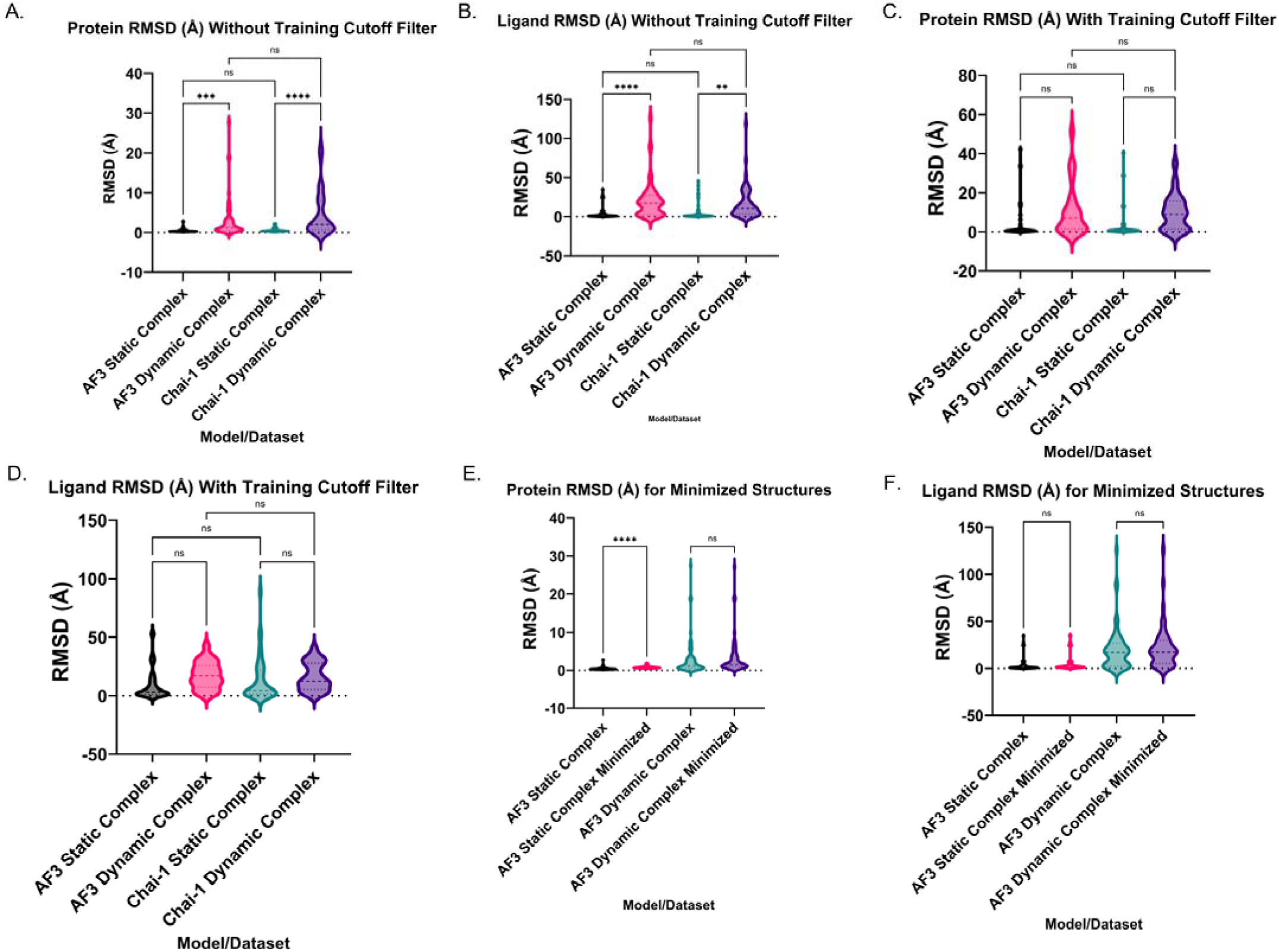
Protein/ligand RMSD between predicted and true structures. A, B: Protein backbone RMSD and ligand RMSD for structures without training cutoff filter (January 2021). C, D: Protein backbone RMSD and ligand RMSD for structures with the training cutoff filter. E, F: Protein backbone RMSD and ligand RMSD for minimized structures before training cutoff using OpenMM’s Minimize() function. (Statistical significance was determined using Brown-Forsythe and Welch ANOVA with Dunnett’s T3 multiple comparison, *p < 0.05, **p < 0.01, ***p < 0.001)

**Figure 2.**
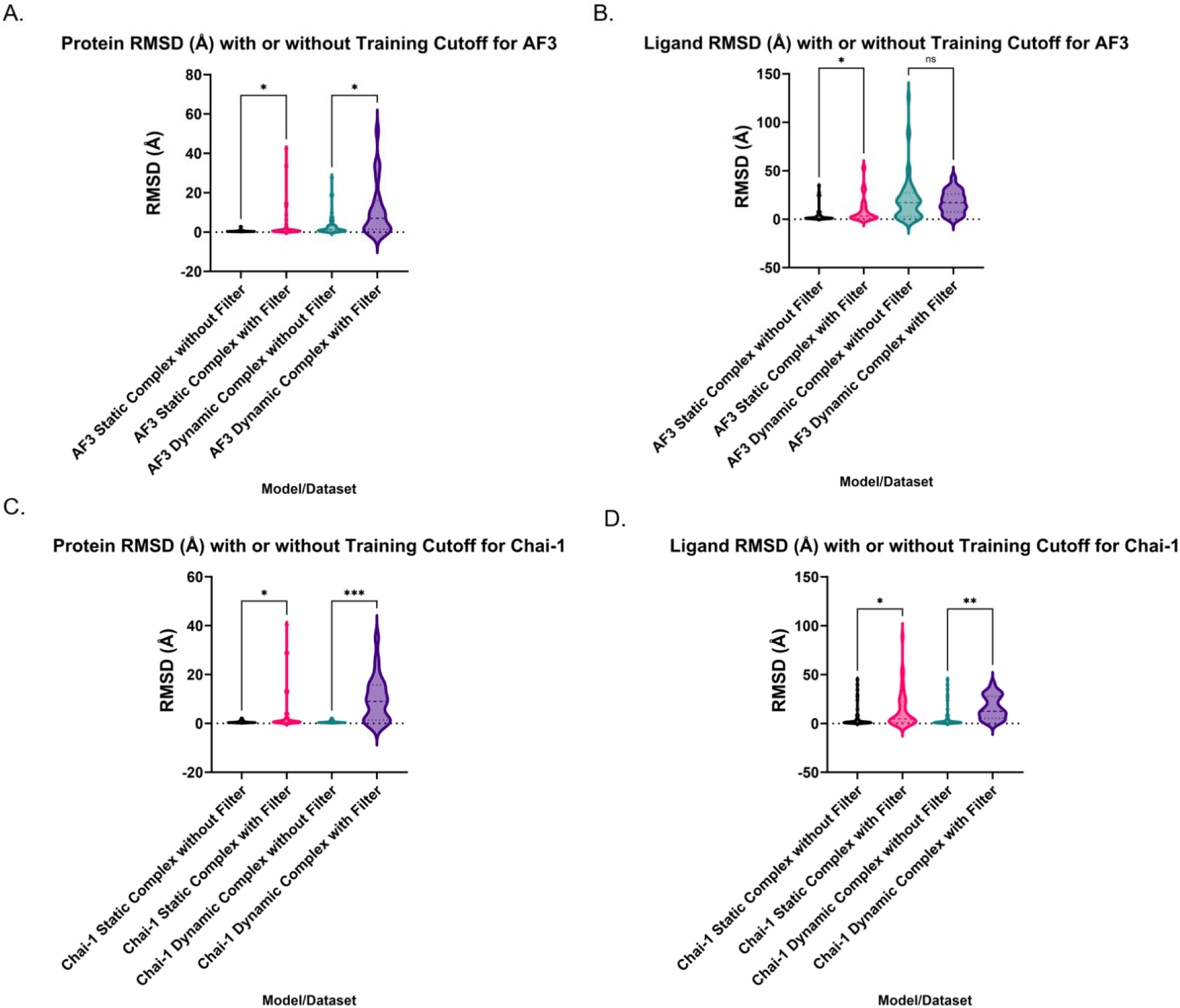
Performance comparison on structures released before and after training cutoff. A, B: Comparison of protein backbone RMSD and ligand RMSD for structures without or with training cutoff filter for AF3. C, D: Comparison of protein backbone RMSD and ligand RMSD for structures without or with training cutoff filter for Chai-1. (Statistical significance was determined using Brown-Forsythe and Welch ANOVA with Dunnett’s T3 multiple comparison, *p < 0.05, **p < 0.01, ***p < 0.001)

We further explored whether energy minimization could enhance prediction quality by applying OpenMM’s Minimize function to optimize atomic positions.^11^ This post-processing step yielded appreciable improvements specifically for protein backbone predictions in static complexes modeled by AF3, suggesting selective benefits of energy refinement (Figure 1E and F).

Beyond geometric alignment, we examined the recovery of specific protein-ligand interactions, which are primarily determined by side-chain conformations and ultimately dictate binding affinity. Our automated analysis using Schrodinger’s Maestro identified comparable interaction recovery rates between AF3 and Chai-1 for structures not filtered by training cutoff date, with both models performing better on Static Complexes than Dynamic Complexes (Figure 3A). This advantage notably disappeared for post-cutoff structures (Figure 3B), providing compelling evidence of training data memorization rather than generalizable learning. Direct comparison of recovery rates confirmed significantly reduced accuracy on novel structures for both models in Static Complex prediction (Figure 3C and D).

**Figure 3.**
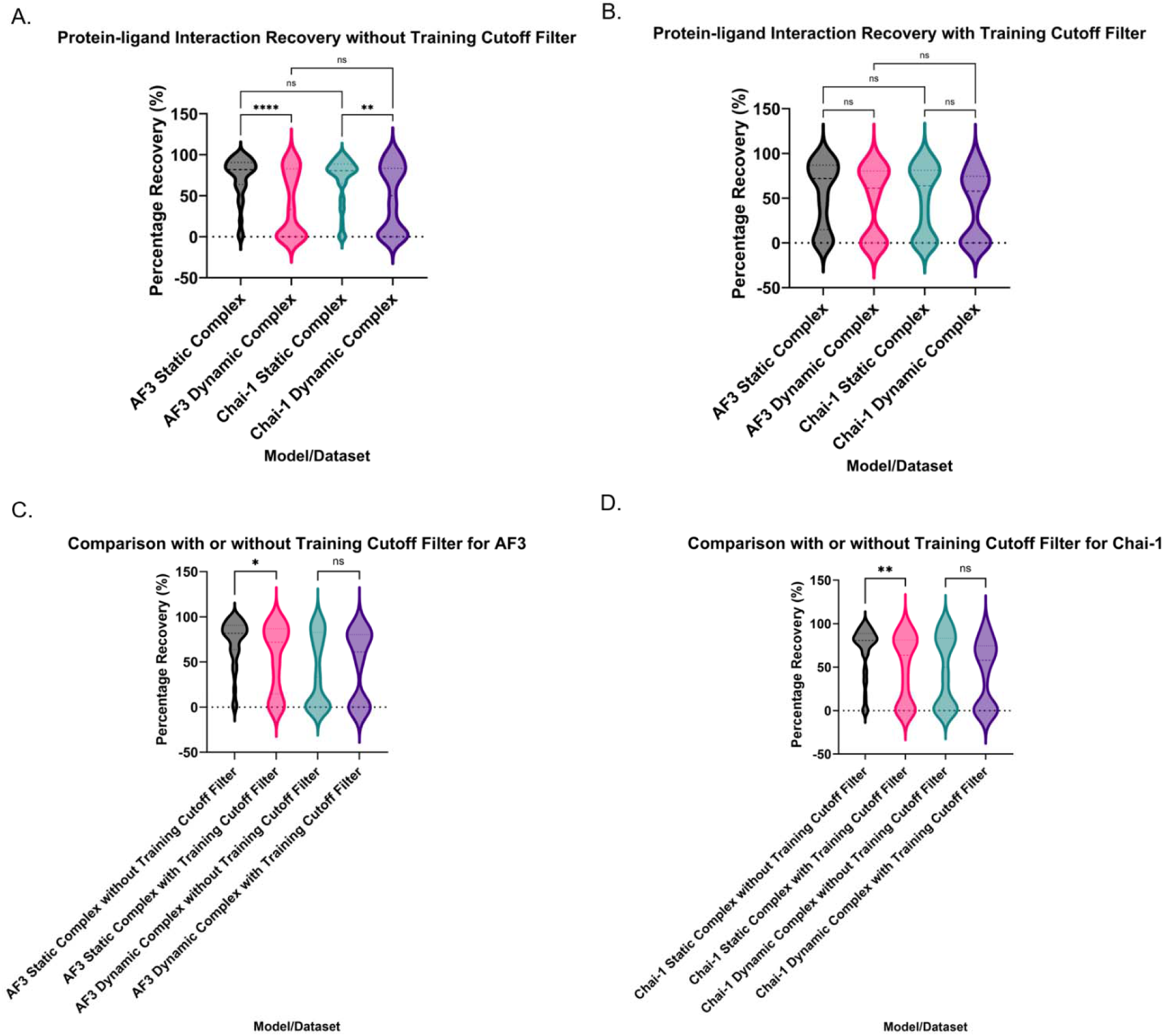
Protein-ligand interaction recovery percentage between predicted and true structures. A, B: Interaction recovery rate without or with training cutoff filter. C, D: Performance comparison for AF3 and Chai-1 between structures without or with training cutoff filter. (Statistical significance was determined using Brown-Forsythe and Welch ANOVA with Dunnett’s T3 multiple comparison, *p < 0.05, **p < 0.01, ***p < 0.001)

To validate our computational findings, six human experts (medicinal chemists) conducted blind manual assessments of randomly selected structures. This human evaluation corroborated our automated analyses, confirming significantly higher success rates for static versus dynamic complexes (Figure 4A). We established strong concordance between human judgment and automated metrics, with successful predictions (defined as >70% interaction recovery) closely aligning with expert assessments (Figure 4B). Confusion matrix (Figure 4C) further demonstrates the strong alignment between automated analyses and human evaluation. Statistical analysis with Chi-Square tests demonstrated highly significant associations between human evaluations and automated measurements for both models (p = 8.15×10⁻ for AF3; p = 3.9×10⁻ for Chai-1).

**Figure 4.**
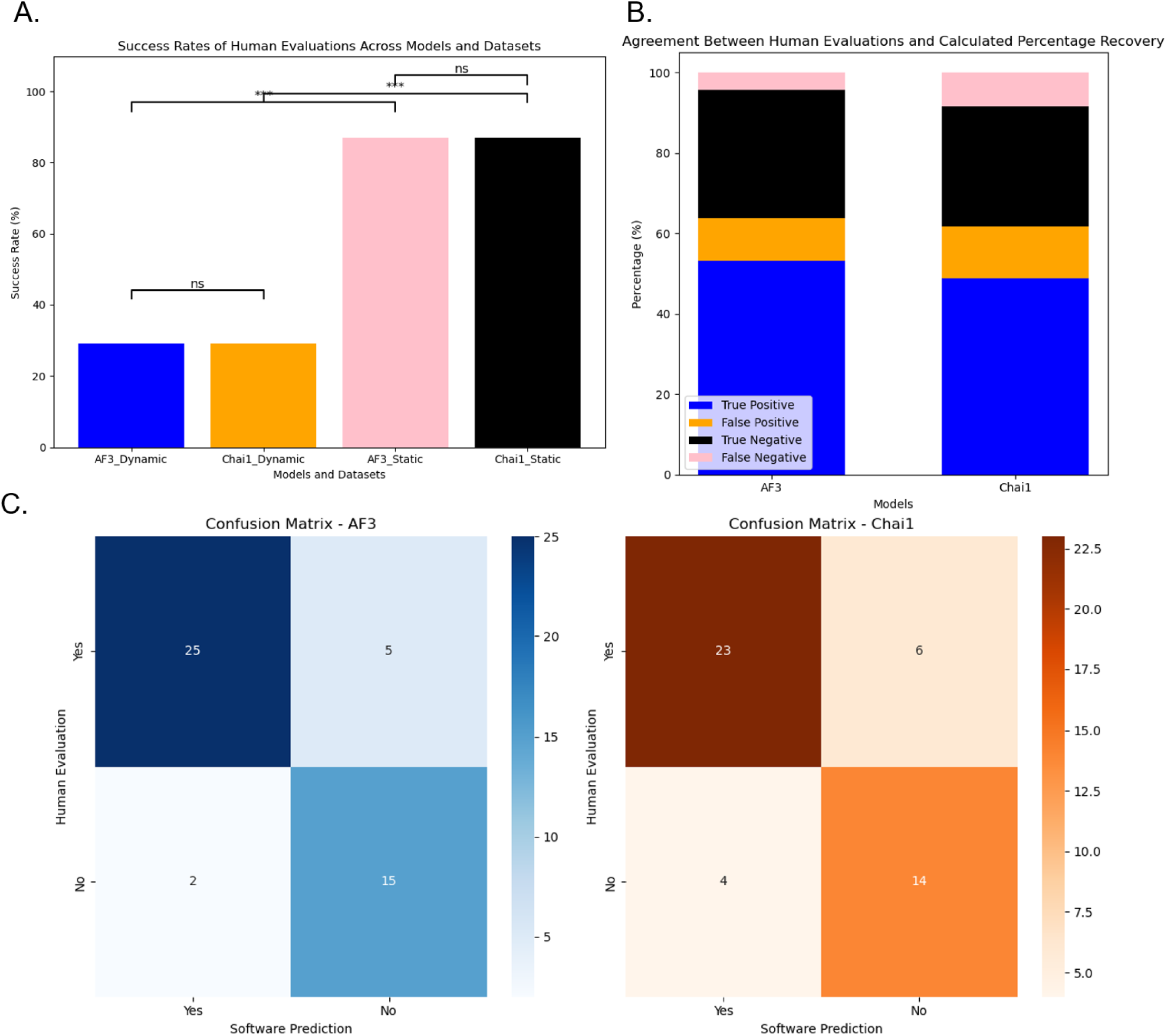
Protein-ligand interaction evaluations from human experts. A: Success rate of AF3 and Chai-1 in different datasets. B: Agreement between human evaluations and calculated percentage recovery. C: Confusion matrix for AF3 and Chai-1 human evaluations and calculated percentage recovery. (Statistical significance was determined using a z-test, *p < 0.05, **p < 0.01, ***p < 0.001)

These findings highlight both the remarkable capabilities and fundamental limitations of current AI prediction models. While they excel at modeling static protein-ligand interactions, they demonstrate substantial difficulties capturing the dynamic conformational changes essential for many drug-target recognition events. The reduced performance on post-cutoff structures reveals concerning dependency on training data memorization rather than physical understanding of molecular recognition principles. This has significant implications for drug discovery applications, suggesting these models may be most reliable for targets with minimal binding-induced conformational ch nges, but require careful validation and potential complementary approaches for dynamic systems. Future architectural improvements should focus on enhancing the representation of protein flexibility and conformational sampling to address these limitations.

### Evaluation: GPCR Conformation Prediction - Conformational Bias in AI Prediction Models

G protein-coupled receptors (GPCRs) represent one of the most pharmaceutically important protein families, accounting for approximately 34% of FDA-approved drugs.^12^ The functional versatility of GPCRs stems from their ability to adopt distinct conformational states—primarily active and inactive—that determine downstream signaling outcomes. Agonists stabilize active conformations, triggering signaling cascades, while antagonists preferentially bind to and stabilize inactive conformations, blocking signaling. However, accurate predictions of GPCR’s state in presence of ligands are still a challenge.^13^

We conducted a focused assessment of AF3’s ability to predict these conformational states using the calcium-sensing receptor (CaSR), a Class C GPCR with well-characterized pharmacology. This target provided an ideal test case due to the recent publication by the Shoichet group of a diverse set of experimentally validated agonists and antagonists.^14^ Our analysis revealed a striking limitation: AF3 consistently predicted active conformations for CaSR regardless of whether the input ligand was an agonist or antagonist (Figure 5A and 5B). Quantitatively, all predicted structures showed significantly lower RMSD values when compared to reference active conformations than when compared to inactive conformations, indicating a strong systematic bias.

**Figure 5.**
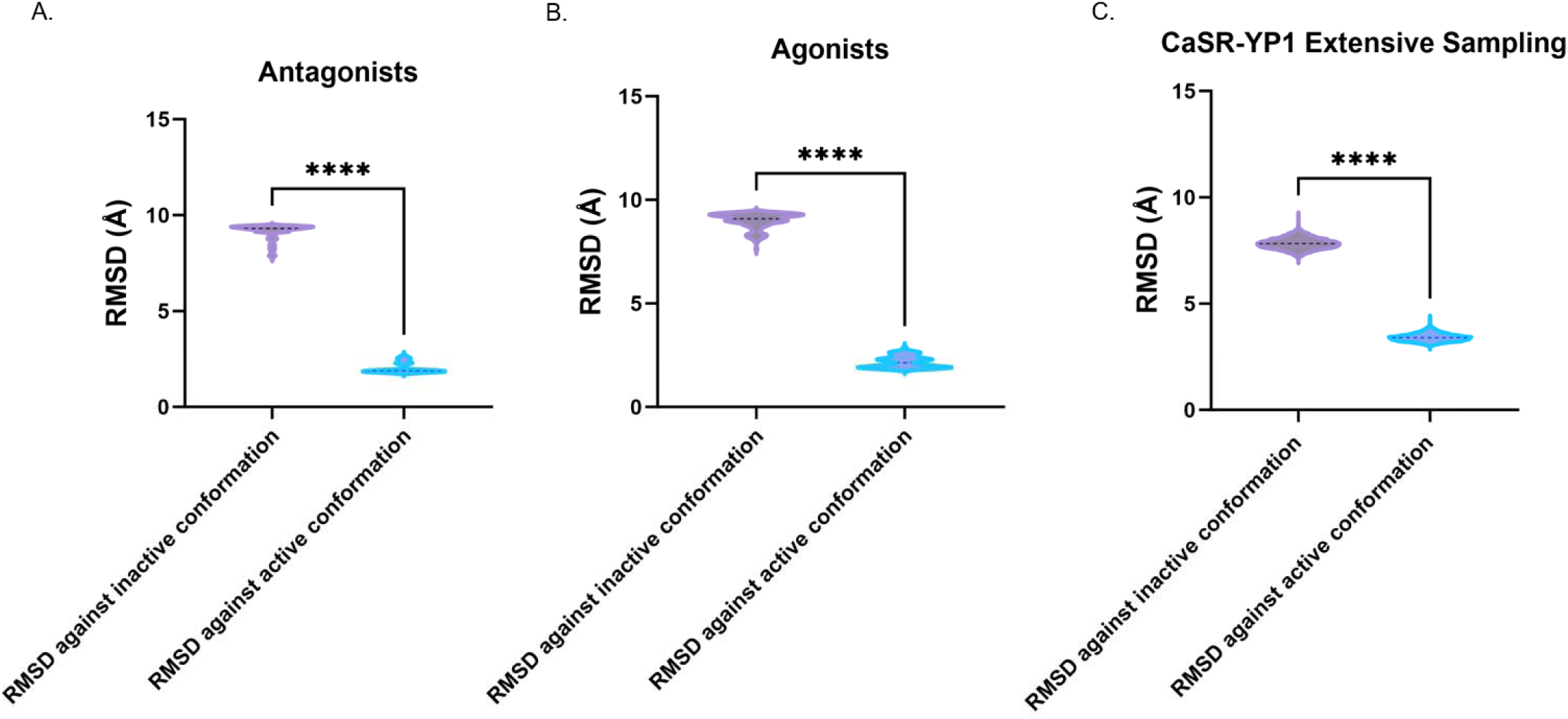
Prediction of CaSR-Compounds Complex. A, B: Protein backbone RMSD of predicted structures agains active conformation and inactive conformation for both agonists and agonists. C: Protein backbone RMSD against activated and inactivated conformation by extensive sampling of CaSR-YP1 complex with AF3. (Statistical significance was determined using paired t-tests, *p < 0.05, **p < 0.01, ***p < 0.001)

To determine whether this conformational bias could be overcome through more extensive computational sampling, we conducted an in-depth analysis using a known antagonist (YP1, PDB ID: 7SIN) that experimentally stabilizes the inactive conformation.^15^ Despite generating approximately 8000 structural predictions using different random seeds across multiple GPUs, AF3 invariably produced active-state conformations (Figure 5C). This consistent failure to predict the correct inactive conformation, even with extensive sampling, points to a fundamental limitation in AF3’s current implementation.

This conformational bias has significant implications for drug discovery applications. The inability to predict antagonist-bound inactive conformations severely limits AF3’s utility for virtual screening campaigns targeting GPCRs, as it would likely miss or mischaracterize antagonists and inverse agonists. More fundamentally, it suggests that AF3’s training methodology may have incorporated an imbalanced representation of GPCR conformational states, potentially reflecting biases in the underlying structural database where active conformations might be overrepresented.

For the broader field of AI-driven structural prediction, this finding highlights the importance of conformational diversity in training datasets and the need for models that can explore alternative conformational states. Future development efforts should focus on implementing enhanced sampling techniques or biasing potentials that can guide the model toward exploring diverse conformational states when appropriate.

### Evaluation: Ternary Complex Prediction Challenges

Ternary complexes—systems involving two proteins connected by a single ligand—represent one of the most challenging frontiers in structural biology and an area of intense pharmaceutical interest. Molecular glues and proteolysis-targeting chimeras (PROTACs) exemplify this class of system, with their therapeutic potential stemming from the induced protein-protein interactions that enable novel biological effects. We evaluated AF3’s capability to predict these complex assemblies using a diverse set of experimentally characterized ternary systems.

Our analysis revealed significant limitations in AF3’s performance on this task. When comparing predicted structures to experimental references, we observed generally poor agreement, with high protein backbone RMSD values as shown in Figure 6A. The complexity of these systems pres nted analytical challenges as well; many predicted structures could not be properly analyzed due to the complicated structures of PROTACs and molecular glues.

**Figure 6.**
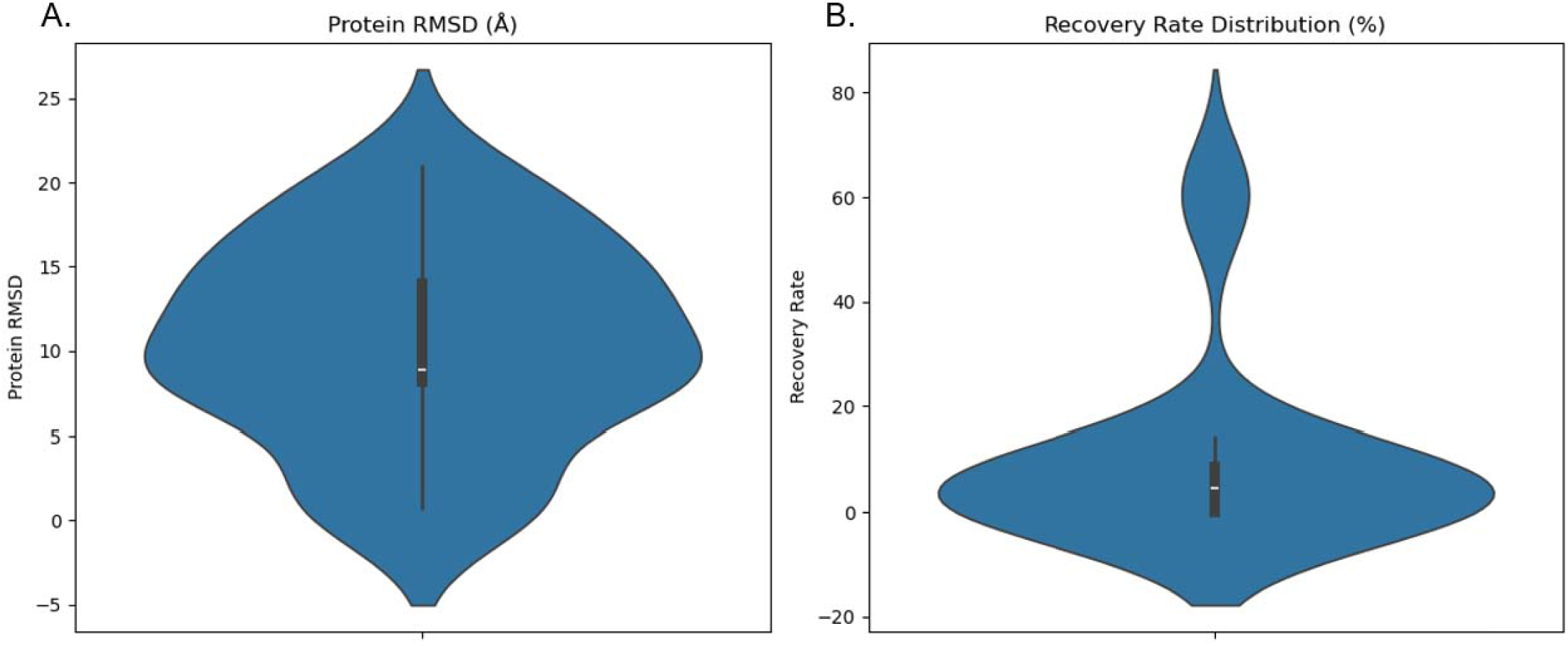
Protein RMSD and PPI recovery rate for ternary systems’ prediction.

Beyond the overall structural alignment, we specifically examined protein-protein interactions (PPIs), which are often central to the therapeutic mechanism of these molecules. By calculating the recovery rate of amino acid-level interactions between the two protein components, we found that AF3 generally failed to accurately recapitulate these critical interfaces (Figure 6B). This shortcoming is particularly significant because the pharmaceutical efficacy of molecular glues and PROTACs fundamentally depends on their ability to induce specific protein-protein contacts.^16^

Interestingly, our analysis revealed some target-dependent variation in performance. For instance, KRAS-related complexes showed relatively higher PPI recovery rates (>70% in some cases), potentially reflecting specific characteristics of these systems where water-mediated interactions play important bridging roles between protein surfaces. This observation suggests that AF3’s performance may vary depending on the specific mechanism of the ternary complex formation and the nature of the interfaces involved.

These findings have important implications for drug discovery efforts focused on molecular glues and PROTACs. While AF3 represents a breakthrough in binary protein-ligand prediction, its current implementation appears inadequate for reliable ternary complex modeling without substantial methodological adaptations. Researchers working in this space should exercise caution when using AF3 for ternary system prediction and should consider alternative specialized approaches or extensive experimental validation.

### Evaluation: Inhibitor Affinity Prediction Capabilities

A critical application of protein-ligand structure prediction in drug discovery is the ability to rank compounds by their binding affinity, guiding lead optimization efforts. While most AF3 benchmarks focus on predicting structures for known binders, we sought to evaluate whether AF3 could effectively discriminate between compounds with varying binding affinities for the same target.

We selected two well-characterized series of soluble epoxide hydrolase (sEH) inhibitors from the CHEMBL database, comprising compounds with a wide range of experimental pIC_50_ values (pIC_50_ equals to the negative of log_10_IC_50_) but sharing common scaffolds.^17,18,19^ This design allowed us to assess AF3’s sensitivity to subtle structural modifications that significantly impact binding affinity while minimizing confounding variables.

Our initial analysis examined the correlation between experimental binding affinities and two metrics provided directly by AF3: the ranking score and the chain_pair_pae (predicted aligned error). According to AF3’s documentation, these metrics might serve as indicators of prediction confidence and potentially correlate with binding strength. However, as shown in Figure 7, neither metric demonstrated meaningful correlation with experimental pIC_50_ values across both inhibitor series. Even compounds with dramatically different binding affinities (spanning several orders of magnitude) received similar AF3 ranking scores and PAE values.

**Figure 7.**
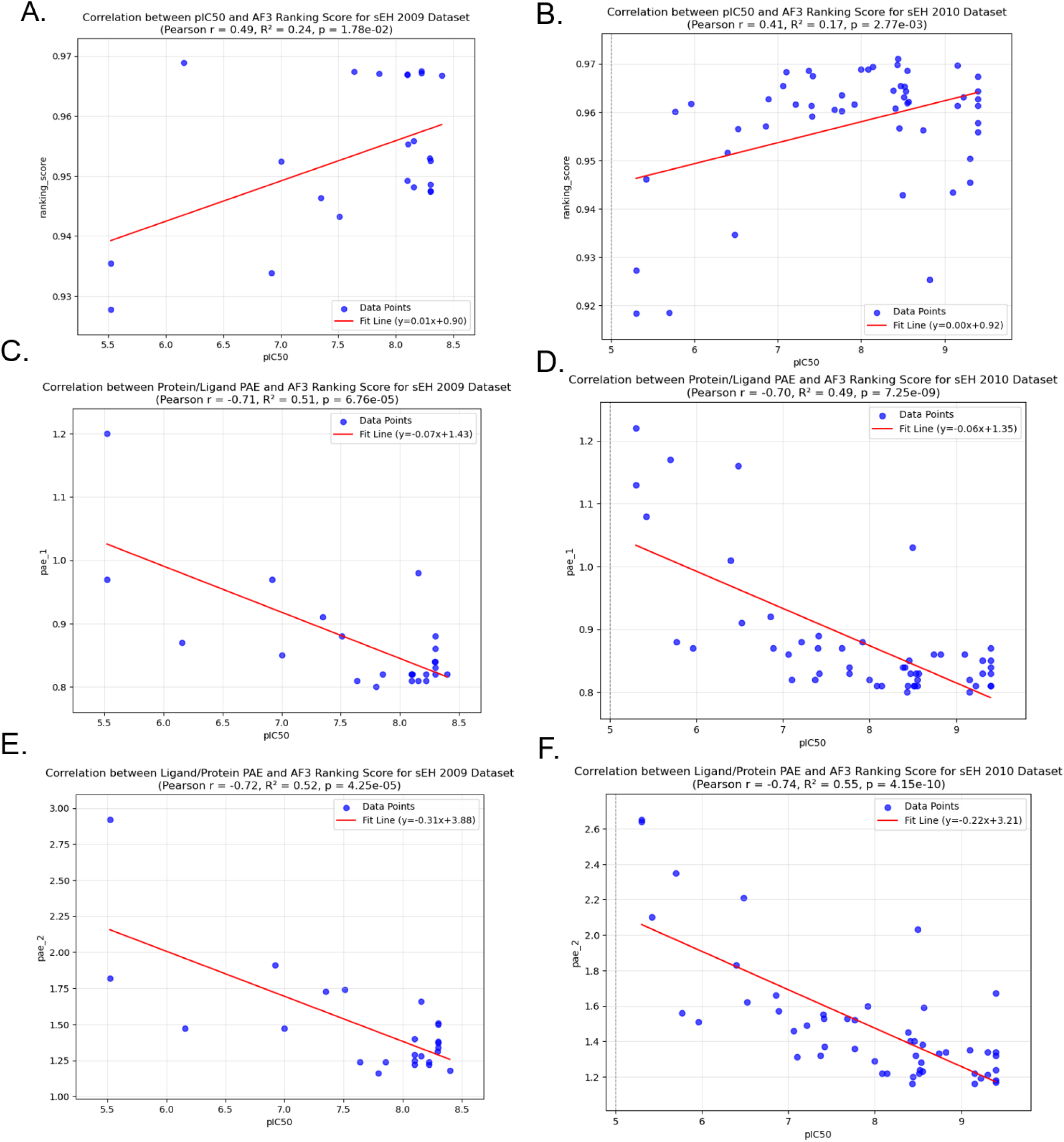
Correlation between pIC50 and AF3 Ranking Score/predicted aligned error for sEH inhibitor series from two publications from 2009 and 2010.

These findings have important implications for drug discovery applications. While AF3 excels at generating high-quality structural models of protein-ligand complexes, it should not be used in isolation for lead optimization campaigns that require accurate affinity ranking. Researchers should consider complementing AF3’s structural predictions with specialized physics-based scoring methods specifically designed for affinity prediction.

The results also highlight the need for continued development of integrated approaches that combine the structural prediction strengths of AI models like AF3 with rigorous energy calculations. Such hybrid methods could potentially transform structure-based drug design by providing both accurate binding modes and reliable affinity estimates from a unified computational pipeline.

### Application Development: Structural Prediction of Chemoproteomics Screening Results

Leveraging our finding that AF3 functions effectively as a true-hit modeler, we applied it to generate structural predictions for protein-ligand interactions identified in a recent large-scale chemoproteomics screening campaign.^20^ The original study reported 47,658 interactions between 407 fragment probes and 2,667 proteins. To maintain computational feasibility while ensuring specificity, we selected a subset of 102 interactions, focusing on proteins with fewer than 11 hits and fragments with fewer than 9 hits. Importantly, we utilized diazirine-alkyne containing probes rather than non-functionalized fragments for our structural modeling, as these bulkier probes provide a more stringent test of binding pocket accommodation—a true binding pocket should have sufficient space for both the probe and its fragment counterpart. Figure 8A illustrates the distribution of Pharmacological Target Development Level (TDL) status across our selected protein hits, providing context for the clinical relevance of these targets. All predicted structures from this analysis are available in the Supplemental Materials and will be accessible through our GitHub repository.

**Figure 8.**
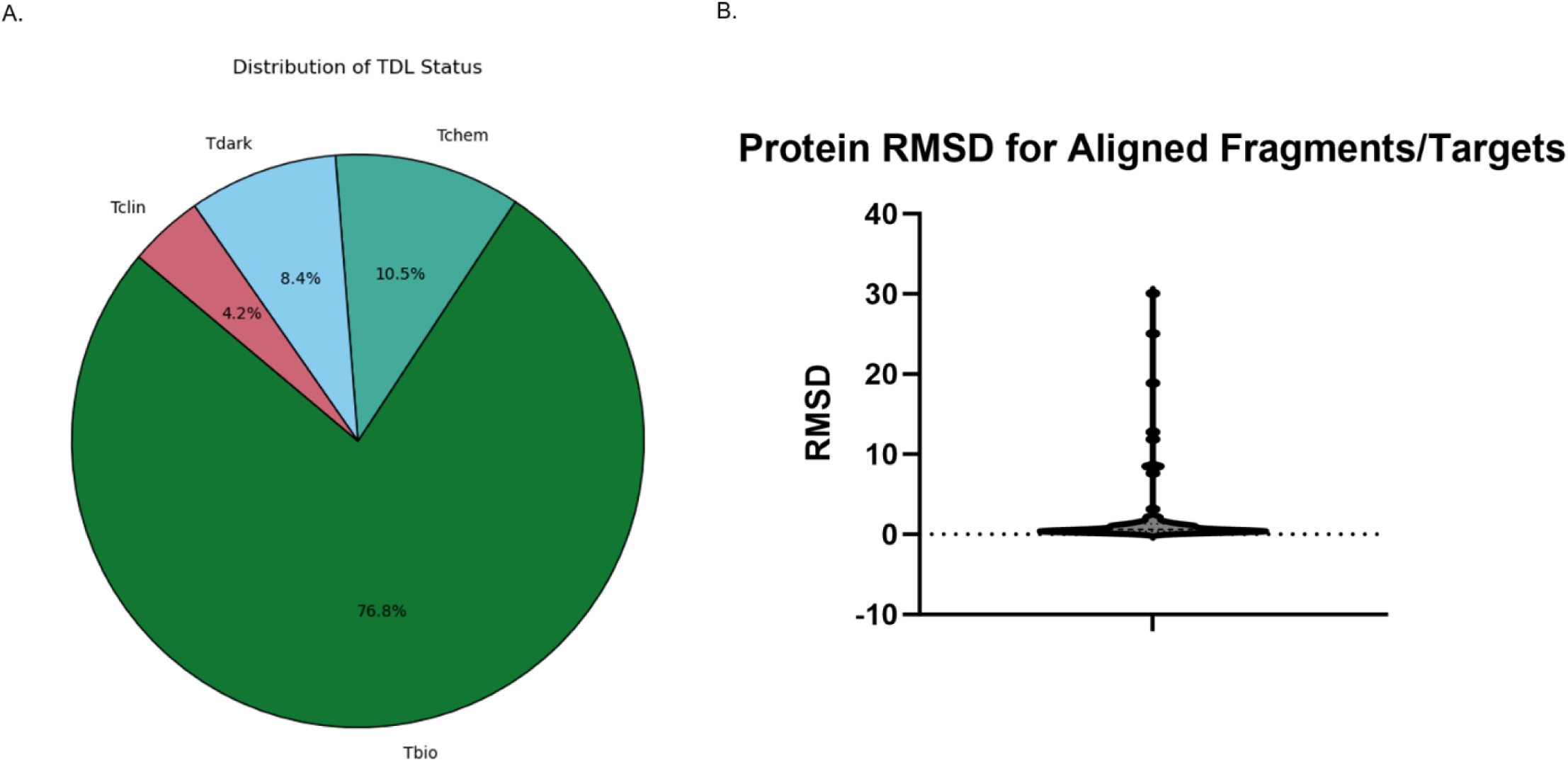
A: Distribution of Pharmacological Target Development Level (TDL) status of selected top 102 hits. B: Protein RMSD for predicted structures with X-ray or Cryo-EM references in RCSB PDB.

To assess the validity of AF3’s structural predictions, we conducted a comprehensive validation using three complementary approaches. First, we compared our generated structures against experimentally determined reference structures available in the PDB. As illustrated in Figure 8B, the majority of our predictions exhibited remarkably low RMSD values when aligned with these experimental structures, providing strong evidence for the reliability of AF3’s predictions in this application context.

Second, we performed targeted validation of binding pocket identification for the subset of targets classified as Tclin and Tchem, representing pharmaceutically relevant proteins with established ligand interactions. Among the 13 targets in these categories, we identified experimental protein-ligand complex structures for 11 targets. Notably, in 9 of these cases, our AF3 predictions correctly identified binding pockets that closely corresponded to those observed experimentally (Table 1). This high concordance rate (82%) confirms AF3’s ability to accurately predict ligand binding sites for known drug targets.

**Table 1.** Pocket validation of Tchem and Tclin targets.

| UniProtID | Gene | TDL_Status | Probe | PDB ID | Ligand Pocket Status |
| --- | --- | --- | --- | --- | --- |
| Q9UBR2 | CTSZ | Tchem | C125 | -- |  |
| Q9NRG4 | SMYD2 | Tchem | C164 | 3S7J | Correct |
| P09417 | QDPR | Tchem | C190, C189 | 1HDR | Correct |
| P51649 | ALDH5A1 | Tclin | C215 | 2W8R | Correct |
| P21281 | ATP6V1B2 | Tchem | C217 | -- |  |
| P05091 | ALDH2 | Tclin | C221 | 3INJ | Correct |
| P07814 | EPRS1 | Tchem | C257 | 4HVC | Correct |
| O15382 | BCAT2 | Tchem | C335 | 1KT8 | Partially Correct |
| Q08752 | PPID | Tchem | C340 | -- |  |
| O95169 | NDUFB8 | Tclin | C360 | -- |  |
| P50579 | METAP2 | Tchem | C416 | 1B6A | Correct |
| P09874 | PARP1 | Tclin | C417 | 4OQA | Correct |
| P07384 | CAPN1 | Tchem | C422 | 8GX3 | Correct |
| P35222 | CTNNB1 | Tchem | C423 | -- |  |

Finally, to prioritize the most reliable structural predictions from our dataset, we implemented a dual-criteria filtration process based on AF3’s ranking score and binding affinity estimation. Specifically, we retained only structures with an AF3 ranking score exceeding 0.7 (indicating high confidence in the overall structural prediction) and a GNINA docking-predicted affinity better than −5 kcal/mol (suggesting strong binding potential).^21^ Figure 9 provides representative examples of structures before and after this filtration process, highlighting the substantial quality difference between models with poor predicted affinity (Figure 9A) versus those with favorable predicted affinity (Figure 9B). This rigorous selection process yielded 21 high-confidence structures that meet both criteria (Chemical structures shown in Table 2, full protein-ligand structures can be found in Supplemental Materials), representing the most promising candidates for further investigation.

**Figure 9.**
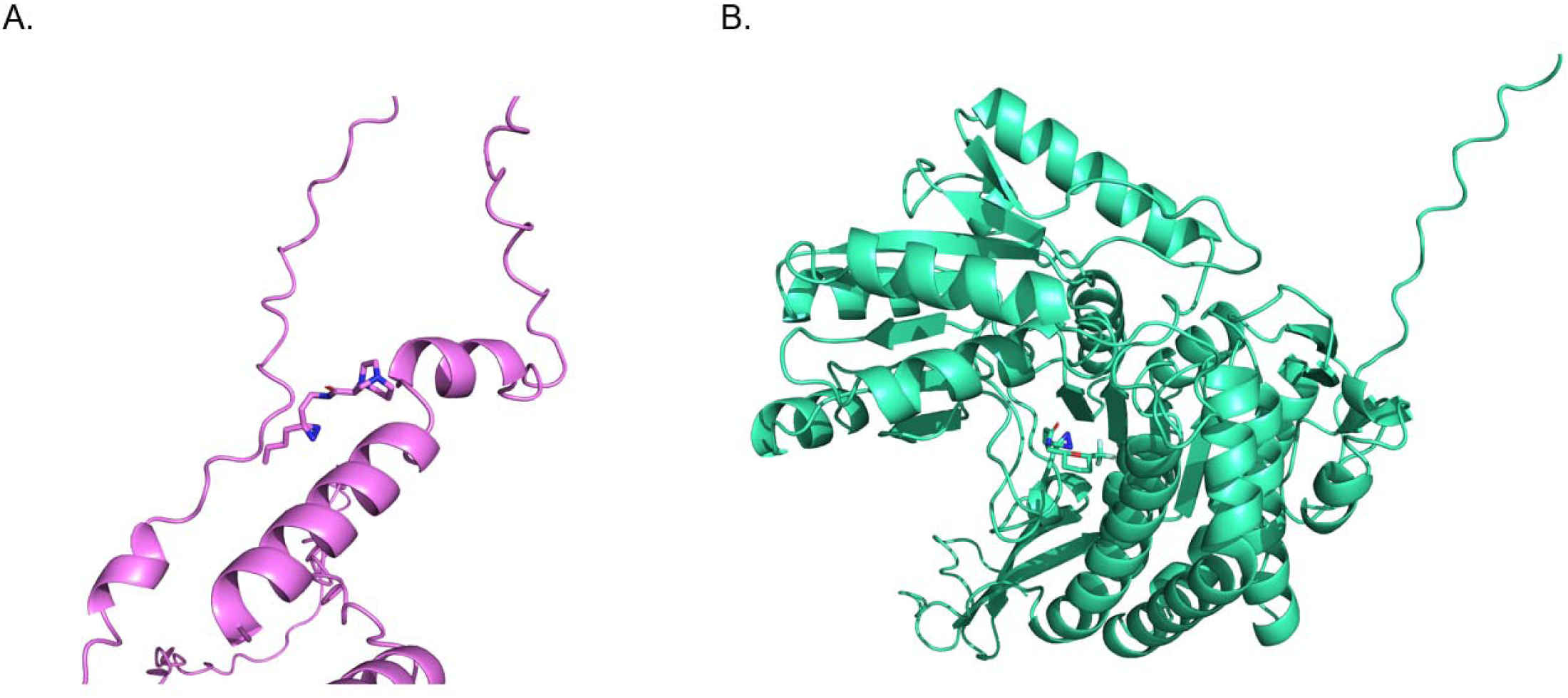
Example structures from unfiltered and filtered structures. A. Q99935(Opiorphin prepropeptide) with fragment C221. Predicted affinity: −0.56 kcal/mol. B: P49419 (Alpha-aminoadipic semialdehyde dehydrogenase) with fragment C360. Predicted affinity: −7.28 kcal/mol.

**Table 2.**
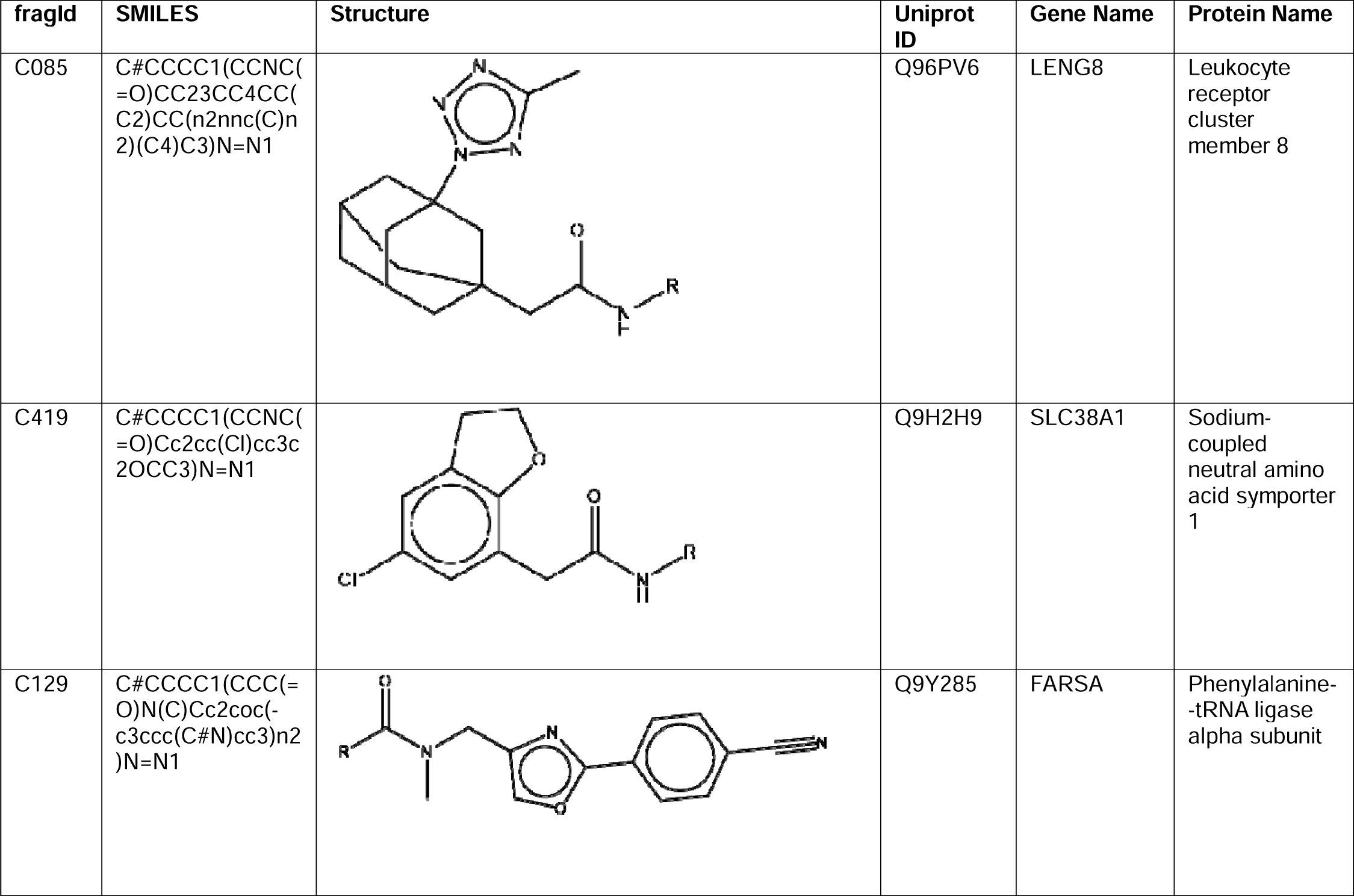
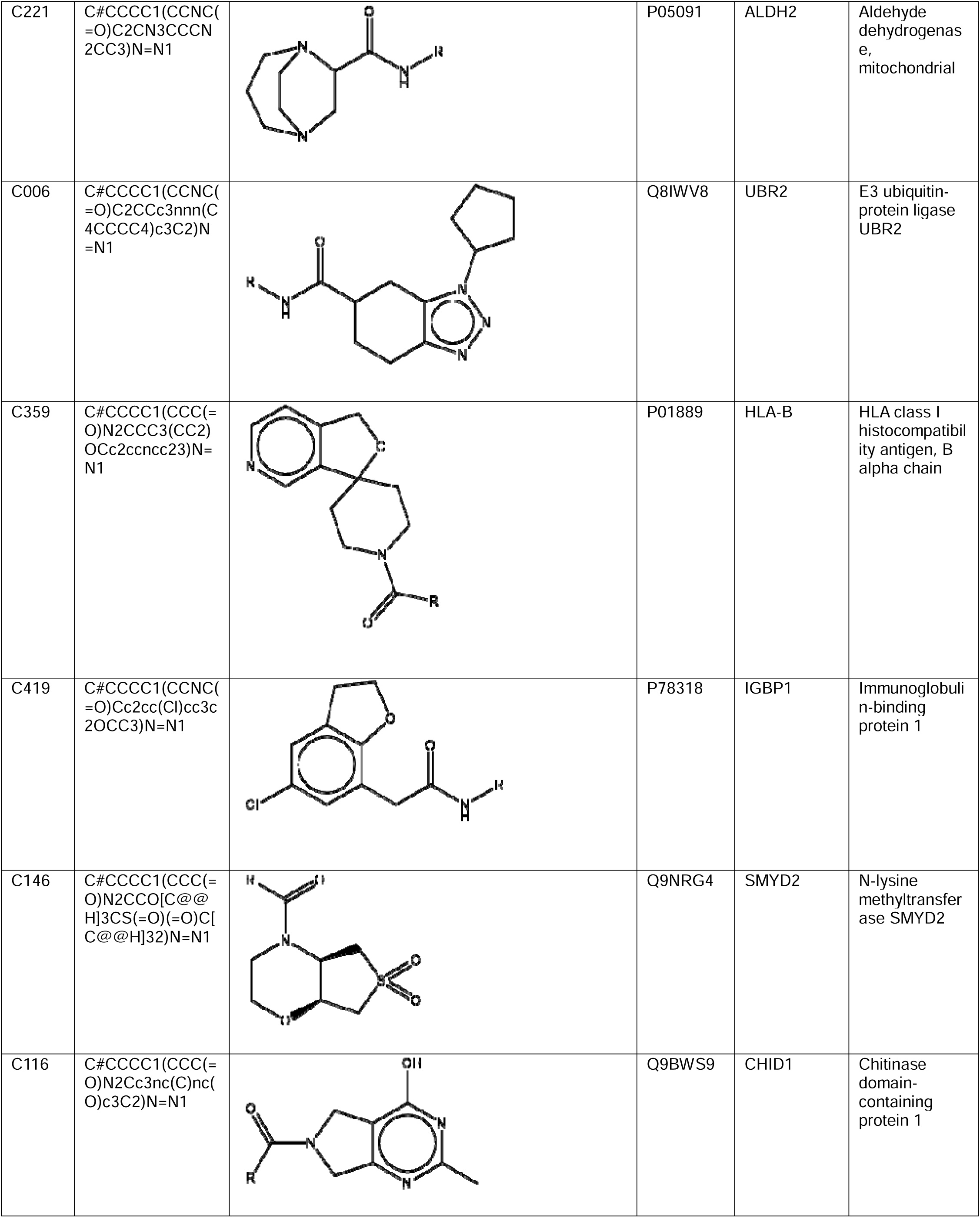
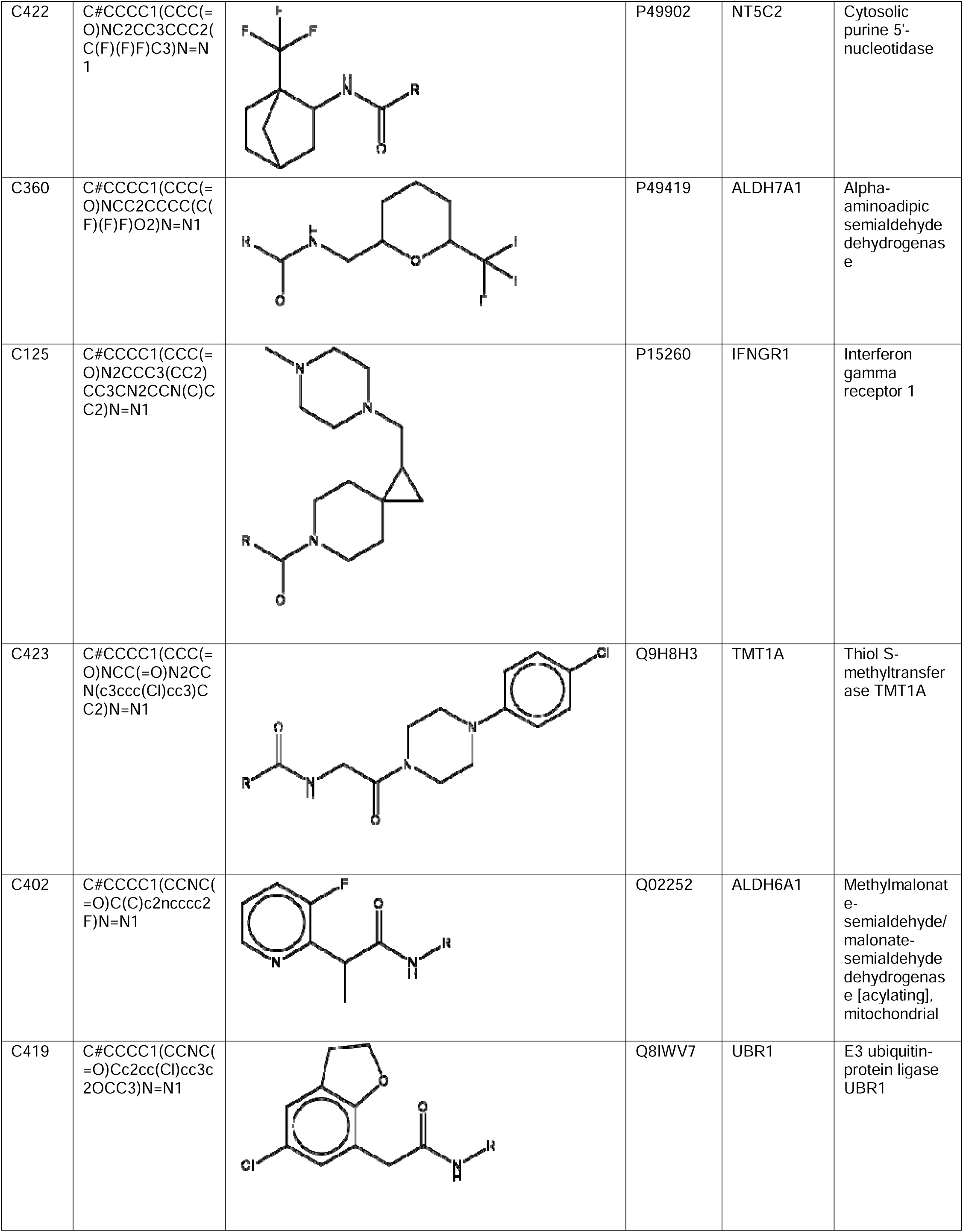
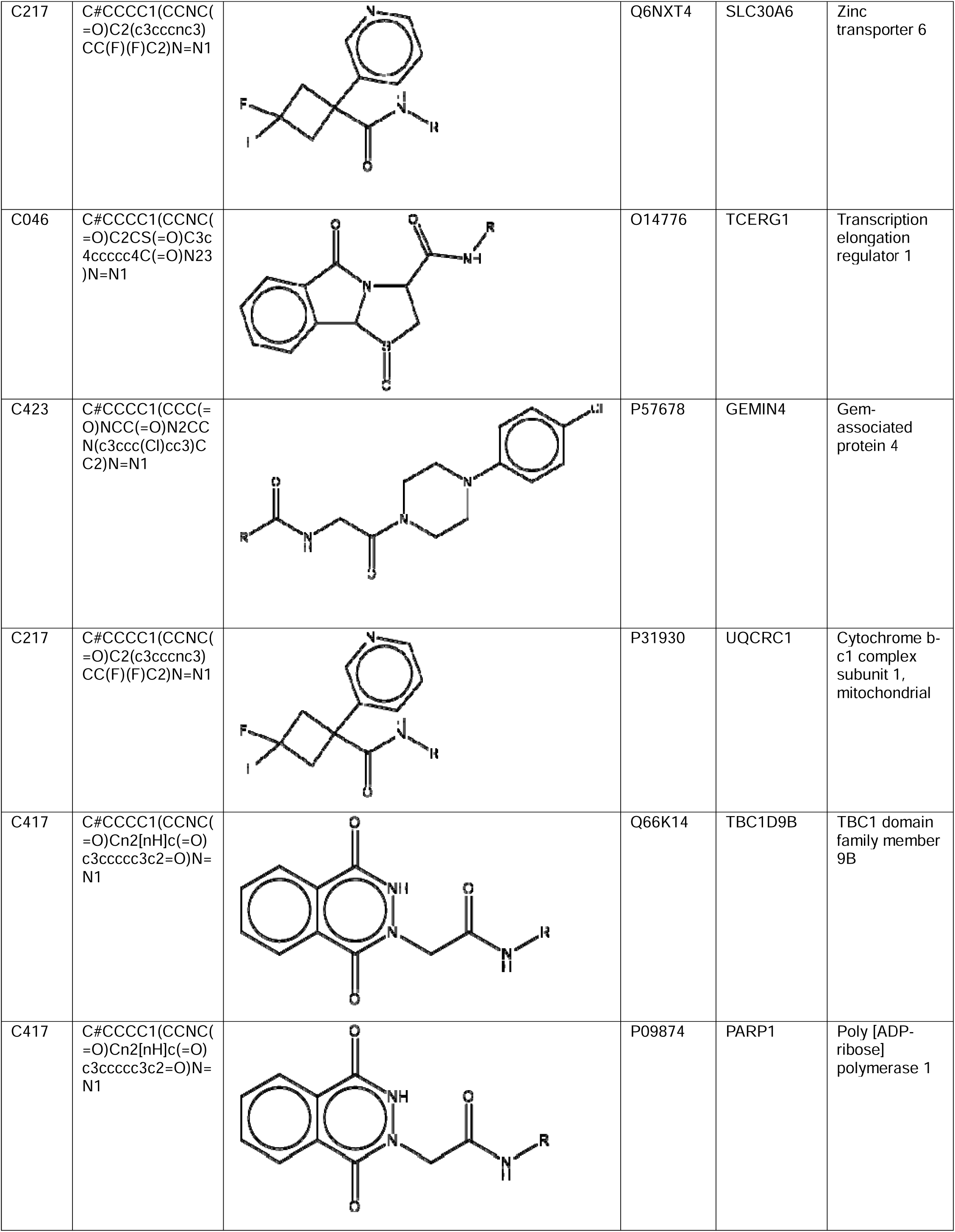
High-confidence fragments selected by AF3 ranking score and GNINA Docking. * R-group: 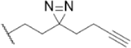

From the 21 high-confidence structures, we selected four protein-fragment pairs for more detailed protein-ligand interaction analysis based on their target status (HLA-B, IFNGR1, UBR1: Tbio, PARP1: Tclin). These proteins are biologically significant or clinically relevant and development of specific ligands will facilitate research on these important targets as chemical probes and create novel therapeutic potential. We draw both 2D and 3D illustrations for these structures in Figure 10 and 11. The result diagrams show that all four fragments are folded and docked into well-defined pockets, and prove that these high-confident predictions can be used for subsequent medicinal chemist y development.

**Figure 10.**
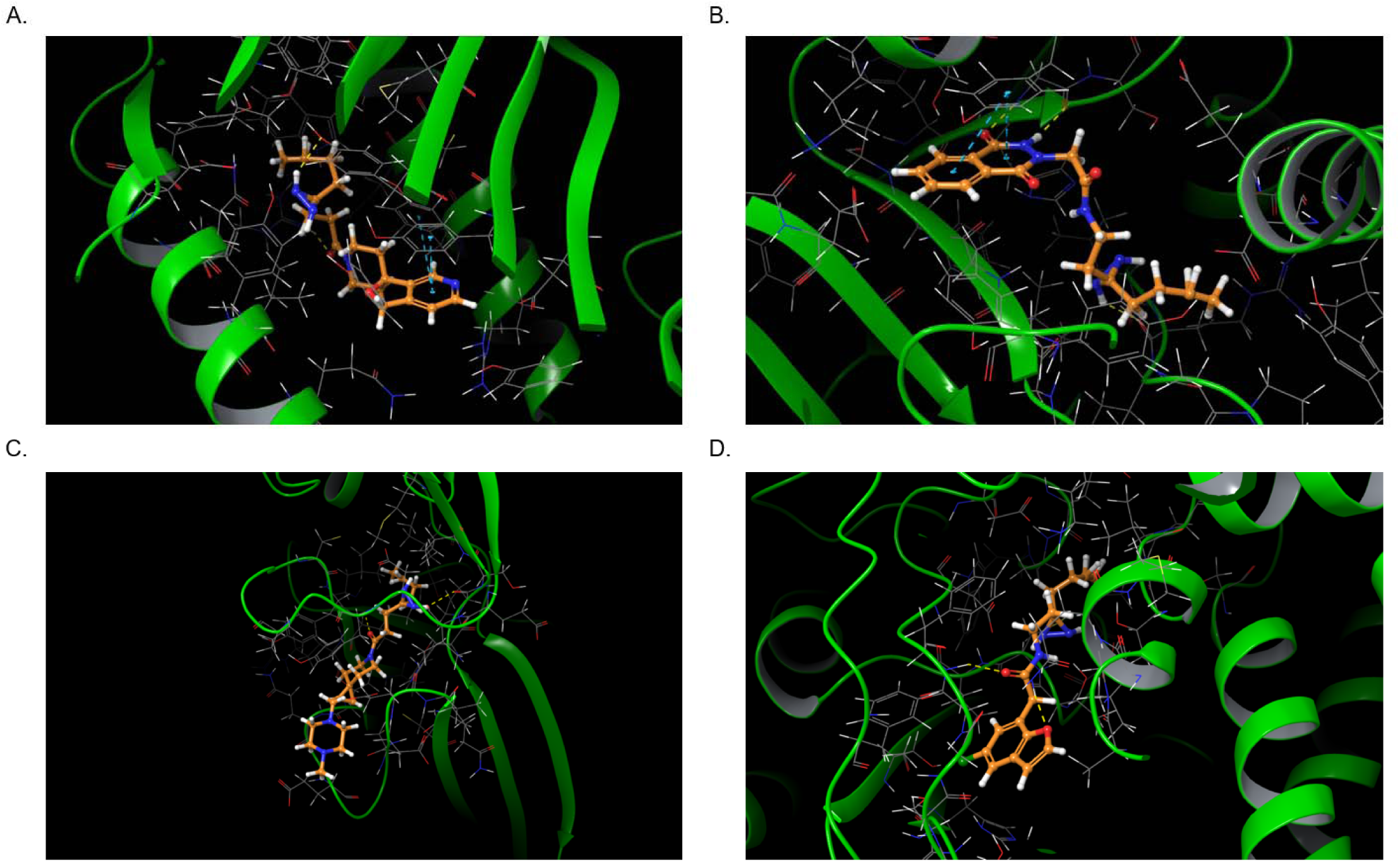
3D protein-ligand interaction diagram for selected fragment-pairs. A: HLA-B with fragment C359. B: PARP1 with fragment C417 C: IFNGR1 with fragment C125 D: UBR1 with fragment C419. Yellow: H-bond, magenta: salt bridges, purple: halogen bond, blue: pi-pi stacking, green: pi-cation

**Figure 11.**
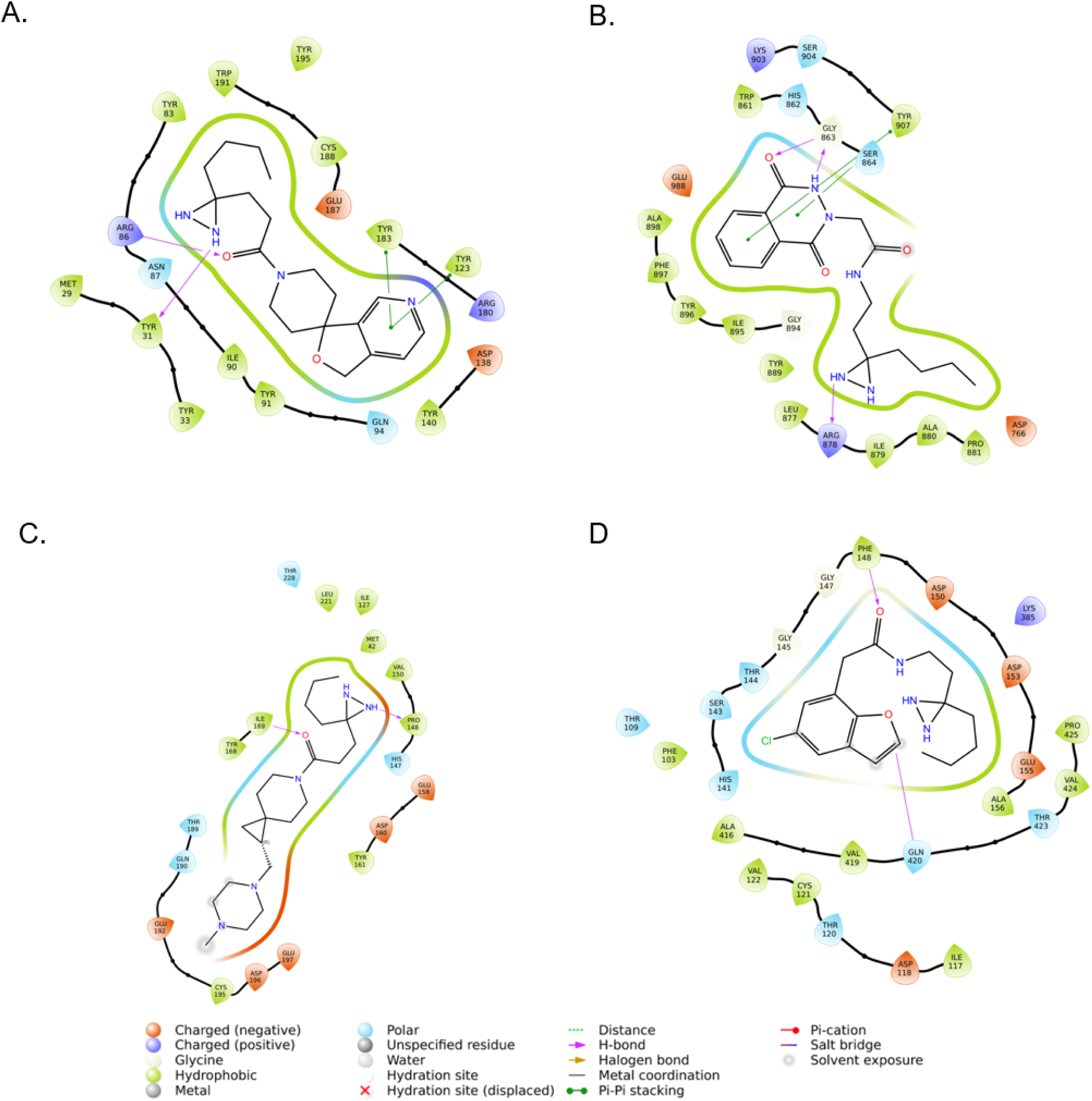
2D protein-ligand interaction diagram for selected fragment-pairs. A: HLA-B with fragment C359. B: PARP1 with fragment C417 C: IFNGR1 with fragment C125 D: UBR1 with fragment C419.

In conclusion, our multi-layered validation approach demonstrates that AF3 can generate reliable protein-ligand structural models from chemoproteomics data, particularly when combined with appropriate filtration strategies. These high-confidence structures provide valuable starting poi ts for structure-based drug design, offering atomic-level insights into protein-ligand interactions that were previously identified only as binary hits in high-throughput screening campaigns.

### Application Development: Prediction of Drug-Resistant Mutants

Drug resistance presents a significant challenge in cancer therapeutics, particularly for targeted kinase inhibitors. To evaluate AF3’s predictive capabilities in this domain, we conducted a systematic analysis of Bruton’s tyrosine kinase (BTK) mutations known to confer resistance to pirtobrutinib, an FDA-approved non-covalent BTK inhibitor, against four non-covalent BTK inhibitors.^22^

Our investigation revealed a notable correlation between AF3’s ranking scores and experimentally determined binding affinities. As shown in the data for pirtobrutinib (Table 3), wild-type BTK exhibits the strongest binding to pirtobrutinib (K_D_ = 0.9 nM) with a corresponding high AF3 ranking score of 0.95. The C481S mutation, which maintains substantial binding (K_D_ = 2.6 nM), shows a similarly high AF3 ranking score of 0.96. More significantly, the highly resistant mutations L528W and A428D, where experimental studies detected no measurable binding, received notably lower AF3 ranking scores of 0.84 and 0.72, respectively. Another NMR based measurement showed that the BTK L528W mutant partially binds to pirtobrutinib at 100 µM concentration^23^, suggesting potentially a very weak binding between pirtobrutinib and the L528W mutant. The A428D mutation completely blocks the kinase pocket entrance. Based on our internal data and a recent study^24^, the A428D mutation is resistant to both kinase inhibitors and degraders^25^. The correlation is not very clear between the experimental *K_D_* and the AF3 ranking scores for ARQ-531, vecabrutinib, and fenebrutinib.

**Table 3.** Predicted CNN affinities, AF3 ranking scores, and experimental dissociation equilibrium constants (K_D_) of BTK variants with BTK inhibitors.

| BTK Variants | Experimental $K_D$ (nM) <sup>a</sup> | AF3 Ranking Score | CNN Affinity |
| --- | --- | --- | --- |
| *Pirtobrutinib* |  |  |  |
| WT | 0.9 | 0.95 | 8.5 |
| A428D | No binding detected | 0.72 | 7.0 |
| M437R | 71 | 0.96 | 8.4 |
| T474I | 14 | 0.93 | 8.2 |
| L528W | No binding detected | 0.82 | 7.6 |
| C481S | 2.6 | 0.96 | 8.5 |
| *ARQ-531* |  |  |  |
| WT | 87 | 0.95 | 8.0 |
| A428D | 2300 | 0.94 | 7.7 |
| M437R | 29 | 0.96 | 8.0 |
| T474I | 8000 | 0.95 | 7.6 |
| L528W | No binding detected | 0.95 | 8.5 |
| C481S | 79 | 0.96 | 7.8 |
| *Vecabrutinib* |  |  |  |
| WT | 0.8 | 0.94 | 9.0 |
| A428D | No binding detected | 0.90 | 8.1 |
| M437R | 1.2 | 0.95 | 8.8 |
| T474I | 14 | 0.95 | 8.5 |
| L528W | 24 | 0.89 | 8.5 |
| C481S | 2.5 | 0.95 | 8.7 |
| *Fenebrutinib* |  |  |  |
| WT | 0.2 | 0.95 | 8.6 |
| A428D | No binding detected | 0.95 | 8.2 |
| M437R | 159 | 0.95 | 8.5 |
| T474I | 2.1 | 0.95 | 8.5 |
| L528W | 1.5 | 0.95 | 8.7 |
| C481S | 5.1 | 0.95 | 8.5 |

a. The dissociation equilibrium constants ( $K_D$ ) of compounds to purified wild-type or mutant BTK protein are determined with the use of surface plasmon resonance technology. Data were adopted from Wang (2022).<sup>22</sup>

This pattern suggests that AF3 effectively captures the structural incompatibilities that underlie resistance mechanisms. Upon detailed structural examination, we observed that these resistance mutations primarily operate through steric hindrance mechanisms. The introduction of bulkier side chains L528W and A428D inside and at the entrance of the binding pocket, respectively, creates spatial constraints that physically prevent optimal inhibitor positioning. AF3 appears particularly adept at recognizing these steric clashes, as evidenced by the progressive decrease in ranking scores that parallels the experimental binding data.

The M437R mutation represents an intermediate case, with reduced but still detectable binding (K_D_ = 71.0 nM) and a high AF3 ranking score of 0.96. This suggests that while AF3 identifies the general binding compatibility, it may not fully capture more subtle electronic or dynamic effects that contribute to moderate resistance phenotypes.

In addition to AF3 ranking scores, we also GNINA docking to dock and score our predicted structures. The difference between CNN Affinities correlates well with the difference between K_D_ within the dataset for the same inhibitors, except for ARQ-531, as shown in Figure 12. This result shows that the docking method could be a useful follow-up tool for ranking affinity for the same compound against different mutations.

**Figure 12.**
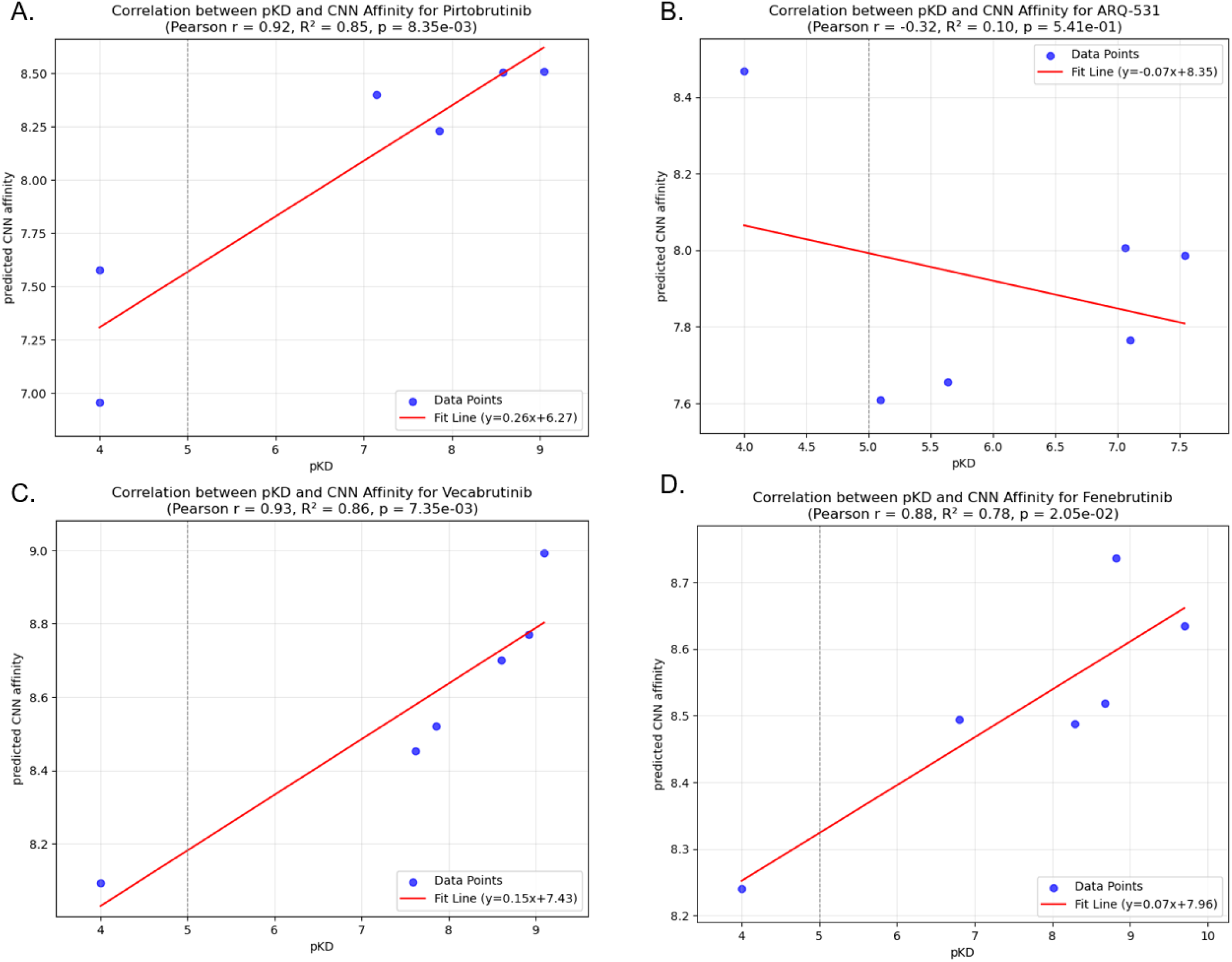
Correlation between GNINA CNN Docking score and pK_D_. “No binding detected” was replaced with 100 μM for plotting purposes.

These findings highlight AF3’s utility in predicting resistance mechanisms driven by structural incompatibilities. While the model may not provide precise quantitative affinity predictions, the ranking scores offer valuable qualitative insights into binding disruptions caused by steric interference. Such predictive capability could prove valuable in anticipating resistance pathways and guiding the development of next-generation inhibitors designed to overcome specific resistance mechanisms.

### Application Development: Kinome Scanning Prediction

Following our insights from BTK inhibitor ranking and mutation studies, we extended our evaluation to assess AF3’s capability to predict kinase selectivity profiles across the human kinome. This application represents a more challenging predictive task that tests whether AF3 can discriminate binding preferences across highly similar yet functionally distinct protein targets.

We developed a computational workflow combining AF3 structural prediction with GNINA docking to simulate kinome profiling experiments. Using data from the Library of Integrated Network-Based Cellular Signatures (LINCS), we selected three well-characterized kinase inhibitors with diverse selectivity profiles: AMG706 (Motesanib), GDC-0941 (Pictilisib), and XMD-1150. For each compound, we generated structural predictions across 379 kinases and compared the computational predictions with experimental binding data.^26,27^

The resulting ROC curves (Figure 13) reveal that our AF3-GNINA pipeline achieved only marginally better predictive performance than random assignment across all three test cases. This poor discriminatory power stands in marked contrast to our findings with BTK resistance mutations, where AF3 demonstrated reasonable ability to identify binding incompatibilities caused by steric clashes.

**Figure 13.**
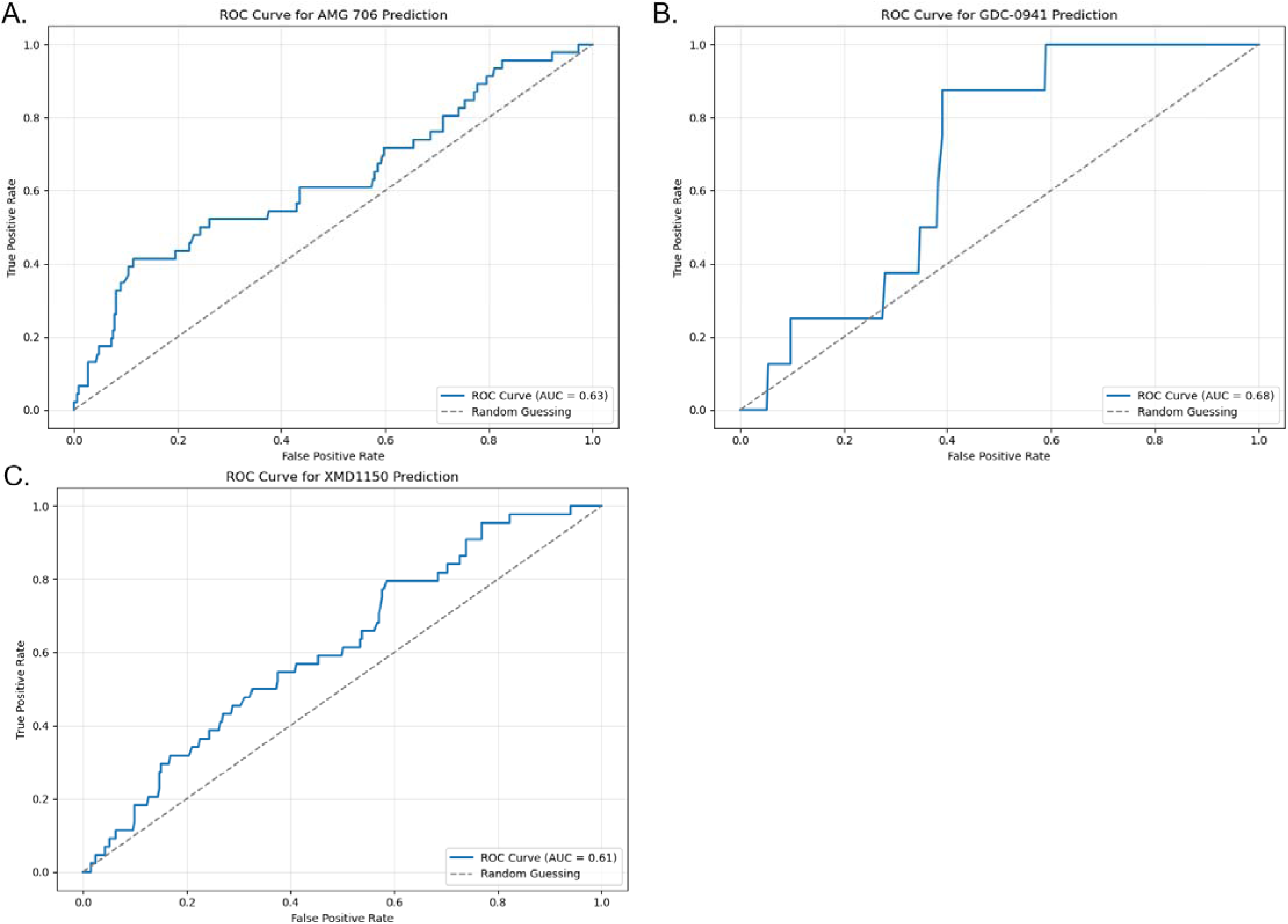
ROC Curves for kinome scanning prediction. A: AMG706, B: GDC-1041, C: XMD1150

This performance discrepancy provides valuable insights into the mechanistic limitations of AF3’s predictive capabilities. While the model appears effective at identifying gross structural incompatibilities (as in the case of BTK mutations that introduce steric hindrance), it struggles to accurately model the more subtle determinants of binding selectivity across the kinome. Kinase selectivity often derives from the absence of specific favorable interactions rather than the presence of outright steric clashes. These missing interactions—whether hydrogen bonds, π-stacking arrangements, or complementary electrostatic surfaces—represent energetic opportunities not realized, rather than explicit structural conflicts.

The juxtaposition of these two findings suggests a fundamental asymmetry in AF3’s predictive capabilities: the model more reliably identifies factors that actively prevent binding (negative determinants) than it recognizes the specific interaction patterns required to enable binding (positive determinants). This asymmetry likely reflects inherent limitations in the model’s training methodology, which may emphasize structural compatibility over the energetic nuances that ultimately determine binding affinity and selectivity.

From a practical standpoint, these findings suggest that AF3 may be more valuable for predicting resistance mechanisms arising from steric conflicts than for prospectively identifying selective compounds across target families. Drug developers should exercise caution when applying AF3 to predict kinase selectivity profiles, as the current implementation appears insufficient for this challenging task without substantial methodological refinement.

## Discussion

In this study, we conducted a comprehensive assessment of AF3 across diverse drug discovery scenarios to systematically evaluate its capabilities and limitations. Our findings reveal a nuanced picture of AF3’s potential applications in pharmaceutical research, with clear performance boundaries that must be considered when integrating this tool into drug discovery workflows.

AF3 demonstrates exceptional performance in predicting static protein-ligand interactions, achieving low RMSD values and high interaction recovery rates for systems that undergo minimal conformational change upon binding. This capability represents a substantial advancement over traditional docking methods, which frequently struggle with accurate side-chain positioning. However, AF3 consistently underperforms when predicting protein-ligand complexes that involve significant conformational changes (RMSD > 5Å between apo and holo states), fundamentally restricting its utility for proteins that undergo induced-fit binding mechanisms or significant domain movements.

The marked decline in performance on post-training cutoff structures suggests concerning memorization effects rather than physical understanding of molecular recognition principles. This raises important questions about the model’s ability to generalize to novel protein-ligand pairs outside its training distribution. Perhaps the most striking is AF3’s systematic bias toward active conformations in GPCR prediction, even when modeling antagonist-bound states that experimentally adopt inactive conformations. This consistent failure persisted despite extensive sampling, indicating a fundamental limitation in the model’s training methodology or representational capacity.

Our evaluation also revealed poor performance on ternary complexes, highlighting significant challenges in modeling the complex interplay between multiple biomolecules, particularly the induced protein-protein interfaces critical for molecular glues and PROTACs. Additionally, AF3’s inability to reliably rank compounds by binding affinity or distinguish active from inactive compounds through its internal metrics (ranking score and PAE) confirms that the model was not designed for energy calculations.

Based on these findings, we propose that AF3’s primary value in drug discovery workflows lies in its capability as a “true-hit binary interaction modeler.” Given a known binding relationship between a protein and ligand, AF3 excels at generating plausible structural models of their interaction, particularly for systems with minimal conformational plasticity. This capability offers significant advantages over traditional docking approaches, as it autonomously identifies the most probable binding location for a ligand based on the global protein structure, demonstrates superior accuracy in predicting side-chain conformations within binding pockets, and provides atomic-level visualization to guide structure-based optimization efforts. Our success in applying this capability to chemoproteomics data demonstrates its practical utility in transforming binary hit information into structurally detailed binding models that can serve as starting points for lead optimization campaigns.

The performance boundaries identified in our study likely stem from fundamental limitations in AF3’s architectural design and training methodology. The consistent failure to predict GPCR inactive conformations suggests insufficient exploration of conformational space during structure generation.

Unlike molecular dynamics approaches that explicitly sample conformational transitions, AF3’s diffusion-based generation may inherently favor the most statistically prevalent conformations in its training dataset. Furthermore, AF3’s architecture appears to lack explicit representation of the energetic factors that determine binding affinity and selectivity. While it can generate structurally plausible conformations, it does not incorporate the physics-based energy functions necessary for accurate ranking of binding interactions. The binary performance disparity between static and dynamic complexes suggests limitations in modeling the conformational ensembles characteristic of many protein-ligand systems, likely reflecting the predominance of static crystallographic structures in training data and the absence of explicit dynamics in the model’s architecture.

Our findings highlight several promising avenues for enhancing the utility of AI-driven structural prediction in drug discovery. The recent advancements demonstrated by YDS PharmaTech’s Ternoplex model for ternary complex prediction provide compelling evidence that incorporating enhanced sampling techniques can overcome some of AF3’s current limitations.^28^ By building biased sampling strategies into AF3-type models, researchers have achieved significantly improved accuracy for challenging ternary systems. This suggests that the core architecture of AF3 captures substantial interaction information but requires methodological refinements to explore diverse conformational states more effectively.

Combining AF3’s structural prediction capabilities with rigorous physics-based approaches could address the energy calculation limitations identified in our study. Hybrid pipelines that leverage AF3 for initial structure generation followed by free energy perturbation methods (such as FEP+) could potentially achieve both structural accuracy and reliable affinity prediction, albeit at increased computational cost. Developing integrated computational workflows that combine AF3 structural prediction, physics-based scoring, and machine learning-driven compound design could enable more efficient closed-loop drug design systems. Such platforms would iteratively generate, evaluate, and refine candidate molecules based on structural insights from AF3 models, potentially accelerating lead optimization cycles.

Future models would benefit from training datasets explicitly enriched for diverse conformational states, particularly for pharmaceutically relevant target classes like GPCRs. Including molecular dynamics trajectories and NMR ensemble structures alongside static crystallographic data could enhance representation of conformational dynamics. Incorporating explicit energy terms, increased emphasis on physical constraints, and dedicated modules for modeling conformational changes could address core limitations in current architectures. Models that combine the generative power of diffusion approaches with the physical realism of energy-based methods represent a promising direction for next-generation structural prediction tools.

Based on our comprehensive evaluation, we recommend tailoring AF3 applications to its strengths while developing complementary approaches for its limitations. For static protein-ligand systems and structure generation from experimentally validated binding data, AF3 provides superior performance to traditional methods. However, for systems involving significant conformational changes, ternary complexes, or affinity-based compound ranking, researchers should consider alternative or hybrid approaches. The integration of AF3 with physics-based scoring functions, enhanced sampling techniques, and specialized models like YDS Ternoplex could address many current limitations while leveraging AF3’s remarkable capabilities for structural prediction.

In conclusion, while AF3 represents a significant advancement in protein-ligand structure prediction, it is not a universal solution yet for all drug discovery challenges. Its optimal utility lies in generating high-quality structural models for known binding pairs, particularly those involving minimal conformational change. Future developments that address its limitations in energy calculation, conformational sampling, and complex system modeling will be crucial for realizing the full potential of AI-driven approaches in drug discovery. The path forward likely involves not just architectural improvements to AF3-type models but their thoughtful integration into multi-component computational pipelines that combine the strengths of AI-based prediction with physics-based simulation and experimental validation.

## Methods and Materials

### Binary protein-ligand benchmark

To compile datasets for this benchmark, we use PLINDER, a curated and annotated RCSB PDB Dataset.^7^ By searching its link table, PDB entries with both apo/holo structures are selected. Holo structures with apo/holo RMSD smaller than 0.5 are compiled into the Static Complex dataset, while holo structures with apo/holo RMSD larger than 5 Å are compiled into the Dynamic Complex dataset. Then, 150 entries from each dataset were randomly selected and used for benchmarking. Additionally, to check models’ performance on structures they never encountered, we selected the structures released after 2021-01-13 and compiled a new set of Static and Dynamic Complex datasets.

To run AlphaFold3 and Chai-1 simulation for this dataset, UniProt IDs were obtained based on PDB ID and the protein chain ID. SMILES for ligands were read from PLINDER dataset files, and sequences for protein were fetched from UniProt Database based on IDs. All the simulations were run on a local H100 GPU server. Then, the predicted structures were aligned and compared with the true PDB structures. Protein backbone RMSD was calculated by PyMol. Ligand mappings were identified by RDKit^29^ and then ligand RMSD was calculated directly with Python. Predicted protein structures were also relaxed using OpenMM Minimize() function^11^, and compared with experimental structures.

To map the recovery of protein-ligand interactions, we first prepared our prediction and true structures using Schrodinger Maestro prepwizard and then used “poseviewer_interactions.py” script from Schrodinger’s Maestro to detect the interactions between proteins and ligands. The detected interactions were compared between predicted structures and ground truth PDB structures, and recovery percentage was calculated based on matched residue and interaction types. For manual inspection of protein-ligand interactions, around 20 structures from both Static Complex and Dynamic Complex were randomly selected and distributed to six researchers from our lab. Structures were imported into Schrodinger Maestro, repaired, and inspected with Ligand Interaction Diagrams. Interactions between real structures and predicted structures were recorded and compared.

### GPCR Benchmark

Agonists and antagonists of CaSR were fetched from GPCR-DB^8^ based on their AC50 and IC50. The sequence of CaSR and SMILES of these ligands was used to run AF3 folding. The results were aligned with both active and inactive conformations of CaSR to determine the state of folded structures. The result structures were then aligned to both activated and inactivated CaSR structures, and RMSD was calculated for each comparison.

### Ternary Complex Benchmark

Thirty-eight ternary structures were selected. The protein sequences and small molecules’ Chemical Compound Dictionary (CCD) codes were obtained directly from RCSB PDB and used for AF3 folding. Protein-ligand interaction and protein RMSD were calculated with similar workflow as mentioned in Binary protein-ligand benchmark. Protein-protein interactions were identified using Schrodinger Maestro’s protein_interaction_analysis.py script and recovery rate were calculated between real and predicted structures.

### Inhibitor Ranking Benchmark

We searched the CHEMBL database for protein sEH using python REST API and identified several publications containing Medicinal Chemistry series for this target. We then manually inspected the structures and identified two series containing large number of compounds with important scaffolds. The protein sequence and compounds SMILES were used to predict protein-ligand complex structure by AF3. We used GNINA for further scoring of our complex.

### Chemproteomic Result Prediction

Protein sequences were fetched from UniProt database and used in AF3 folding along with ligand Native SMILES. For general structural validation, we used UniProt rest API to query all the PDB X-ray and Cryo-EM structures based on the UniProt ID. The structures with the best resolutions were used to align with predicted structures, and RMSD was calculated with PyMol. For pocket validation with Tclin and Tchem, we manually searched RCSB PDB with the uniport IDs for protein-ligand complex structures. To filter the predicted structures, we selected structures with ranking score > 0.7 and scored them by GNINA docking.

### Drug Resistance Mutants Prediction

BTK sequences with mutations are used to fold with compounds’ SMILES. Then, the predicted complex structures were docked and scored using GNINA docking.

### Kinome Scanning Prediction

379 kinase sequences were fetched from UniProt based on their ID and used along with compound SMILES in AF3 folding. After folding, docking scores were calculated by GNINA docking.

### Data and Code Availability

All the input files, predicted structures, raw data for each table and codes for all the analysis will be available on GitHub.

## Notes

### Competing Interest Statement

J.W. is the co-founder of Chemical Biology Probes LLC. J. W. has stock ownership in CoRegen Inc and serves as a consultant for this company. J.W. is the co-founder of Fortitude Biomedicines, Inc. and holds equity interest in this company.

## References

1. Huang SY, Zou X. Advances and challenges in protein-ligand docking. Int J Mol Sci. 2010;11(8):3016–3034. Published 2010 Aug 18. doi:10.3390/ijms11083016

2. Paggi JM, Belk JA, Hollingsworth SA, et al. Leveraging nonstructural data to predict structures and affinities of protein-ligand complexes. Proc Natl Acad Sci U S A. 2021;118(51):e2112621118. doi:10.1073/pnas.2112621118

3. Jumper J, Evans R, Pritzel A, et al. Highly accurate protein structure prediction with AlphaFold. Nature. 2021;596(7873):583–589. doi:10.1038/s41586-021-03819-2

4. Bryant P, Pozzati G, Elofsson A. Improved prediction of protein-protein interactions using AlphaFold2 [published correction appears in Nat Commun. 2022 Mar 24;13(1):1694. doi: 10.1038/s41467-022-29480-5]. *Nat Commun*. 2022;13(1):1265. Published 2022 Mar 10. doi: http://10.1038/s41467-022-28865-w

5. Lyu J, Kapolka N, Gumpper R, et al. AlphaFold2 structures guide prospective ligand discovery. Science. 2024;384(6702):eadn6354. doi:10.1126/science.adn6354

6. Abramson J, Adler J, Dunger J, et al. Accurate structure prediction of biomolecular interactions with AlphaFold 3. Nature. 2024;630(8016):493–500. doi:10.1038/s41586-024-07487-w

7. Janani Durairaj, Yusuf Adeshina, Zhonglin Cao, Xuejin Zhang, Vladas Oleinikovas, Thomas Duignan, Zachary McClure, Xavier Robin, Gabriel Studer, Daniel Kovtun, Emanuele Rossi, Guoqing Zhou, Srimukh Veccham, Clemens Isert, Yuxing Peng, Prabindh Sundareson, Mehmet Akdel, Gabriele Corso, Hannes Stärk, Gerardo Tauriello, Zachary Carpenter, Michael Bronstein, Emine Kucukbenli, Torsten Schwede, Luca Naef, “PLINDER: The protein-ligand interactions dataset and evaluation resource”, *bioRxiv* July 2024; doi: 10.1101/2024.07.17.603955

8. Pándy-Szekeres G, Munk C, Tsonkov TM, Mordalski S, Harpsøe K, Hauser AS, Bojarski AJ, Gloriam DE. GPCRdb in 2018: adding GPCR structure models and ligands. 2017, *Nucleic Acids Res*., Nov 16. 10.1093/nar/gkx1109

9. Chai Discovery team, Boitreaud J, Dent J, McPartlon M, Meier J, Reis V, Rogozhonikov A, Wu K. Chai-1: Decoding the molecular interactions of life.

10. Raisinghani N, Alshahrani M, Gupta G, et al. Probing Functional Allosteric States and Conformational Ensembles of the Allosteric Protein Kinase States and Mutants: Atomistic Modeling and Comparative Analysis of AlphaFold2, OmegaFold, and AlphaFlow Approaches and Adaptations. J Phys Chem B. 2024;128(45):11088–11107. doi:10.1021/acs.jpcb.4c04985

11. P. Eastman, M. S. Friedrichs, J. D. Chodera, R. J. Radmer, C. M. Bruns, J. P. Ku, K. A. Beauchamp, T. J. Lane, L.-P. Wang, D. Shukla, T. Tye, M. Houston, T. Stich, C. Klein, M. R. Shirts, and V. S. Pande. “OpenMM 4: A Reusable, Extensible, Hardware Independent Library for High Performance Molecular Simulation*.”* J. Chem. Theor. Comput. 9(1): 461–469. (2013).

12. Hauser AS, Attwood MM, Rask-Andersen M, Schiöth HB, Gloriam DE. Trends in GPCR drug discovery: new agents, targets and indications. Nat Rev Drug Discov. 2017;16(12):829–842. doi:10.1038/nrd.2017.178

13. Bartuzi D, Kaczor AA, Targowska-Duda KM, Matosiuk D. Recent Advances and Applications of Molecular Docking to G Protein-Coupled Receptors. Molecules. 2017; 22(2):340. 10.3390/molecules22020340

14. Liu F, Wu CG, Tu CL, et al. Large library docking identifies positive allosteric modulators of the calcium-sensing receptor. Science. 2024;385(6715):eado1868. doi:10.1126/science.ado1868

15. Park J, Zuo H, Frangaj A, et al. Symmetric activation and modulation of the human calcium-sensing receptor. Proc Natl Acad Sci U S A. 2021;118(51):e2115849118. doi:10.1073/pnas.2115849118

16. Riching KM, Caine EA, Urh M, Daniels DL. The importance of cellular degradation kinetics for understanding mechanisms in targeted protein degradation. Chem Soc Rev. 2022;51(14):6210–6221. Published 2022 Jul 18. doi:10.1039/d2cs00339b

17. Zdrazil B, Felix E, Hunter F, et al. The ChEMBL Database in 2023: a drug discovery platform spanning multiple bioactivity data types and time periods. Nucleic Acids Res. 2024;52(D1):D1180–D1192. doi:10.1093/nar/gkad1004

18. Rose TE, Morisseau C, Liu JY, et al. 1-Aryl-3-(1-acylpiperidin-4-yl)urea inhibitors of human and murine soluble epoxide hydrolase: structure-activity relationships, pharmacokinetics, and reduction of inflammatory pain. J Med Chem. 2010;53(19):7067–7075. doi:10.1021/jm100691c

19. Eldrup AB, Soleymanzadeh F, Taylor SJ, et al. Structure-based optimization of arylamides as inhibitors of soluble epoxide hydrolase. J Med Chem. 2009;52(19):5880–5895. doi:10.1021/jm9005302

20. Offensperger F, Tin G, Duran-Frigola M, et al. Large-scale chemoproteomics expedites ligand discovery and predicts ligand behavior in cells. Science. 2024;384(6694):eadk5864. doi:10.1126/science.adk5864

21. Ragoza M, Hochuli J, Idrobo E, Sunseri J, Koes DR. Protein-Ligand Scoring with Convolutional Neural Networks. J Chem Inf Model. 2017;57(4):942–957. doi:10.1021/acs.jcim.6b00740

22. Wang E, Mi X, Thompson MC, et al. Mechanisms of Resistance to Noncovalent Bruton’s Tyrosine Kinase Inhibitors. N Engl J Med. 2022;386(8):735–743. doi:10.1056/NEJMoa2114110

23. Joseph RE, Wales TE, Jayne S, et al. Impact of the clinically approved BTK inhibitors on the conformation of full-length BTK and analysis of the development of BTK resistance mutations in chronic lymphocytic leukemia. Preprint. bioRxiv. 2024;2023.12.18.572223. Published 2024 Oct 10. doi:10.1101/2023.12.18.572223

24. Wong RL, Choi MY, Wang HY, Kipps TJ. Mutation in Bruton Tyrosine Kinase (BTK) A428D confers resistance To BTK-degrader therapy in chronic lymphocytic leukemia. Leukemia. 2024;38(8):1818–1821. doi:10.1038/s41375-024-02317-4

25. Tam CS, Balendran S, Blombery P. Novel mechanisms of resistance in CLL: variant BTK mutations in second-generation and noncovalent BTK inhibitors. Blood. 2025;145(10):1005–1009. doi:10.1182/blood.2024026672

26. Keenan AB, Jenkins SL, Jagodnik KM, et al. The Library of Integrated Network-Based Cellular Signatures NIH Program: System-Level Cataloging of Human Cells Response to Perturbations. Cell Syst. 2018;6(1):13–24. doi:10.1016/j.cels.2017.11.001

27. Davis MI, Hunt JP, Herrgard S, et al. Comprehensive analysis of kinase inhibitor selectivity. Nat Biotechnol. 2011;29(11):1046–1051. Published 2011 Oct 30. doi:10.1038/nbt.1990

28. The Statistical Mechanics Inference Team, Che X. YDS-Ternoplex: Surpassing AlphaFold 3-Type Models for Molecular Glue-Mediated Ternary Complex Prediction. *bioRxiv*. Published December 23, 2024. doi:10.1101/2024.12.23.630090

29. RDKit: Open-source cheminformatics. https://www.rdkit.org

